# Seasonal and environmental factors contribute to the variation in the gut microbiome: a large-scale study of a small bird

**DOI:** 10.1101/2023.12.12.571395

**Authors:** Martta Liukkonen, Jaime Muriel, Jesús Martínez-Padilla, Andreas Nord, Veli-Matti Pakanen, Balázs Rosivall, Vallo Tilgar, Kees van Oers, Kirsten Grond, Suvi Ruuskanen

## Abstract

Environmental variation can shape the gut microbiome, but majority of studies use captive-bred species, while data on large-scale variation in the gut microbiome and the associated environmental factors is lacking. Furthermore, previous studies have limited taxonomical coverage, and for example knowledge about avian gut microbiomes is still scarce. We investigated large-scale environmental variation in the gut microbiome of wild adult great tits across the species’ European distribution range. Our results show that gut microbiome diversity is higher during winter and that there are compositional differences between winter and summer gut microbiomes. During winter, individuals inhabiting mixed forest habitat show higher gut microbiome diversity, whereas there was no similar association during summer. Also, temperature was found to be a small contributor to compositional differences in the gut microbiome. We did not find significant differences in the gut microbiome among populations, nor any association between latitude, rainfall, and the gut microbiome. The results suggest that there is a seasonal change in wild avian gut microbiomes, but that there are still many unknown factors that shape the gut microbiome of wild bird populations.

## INTRODUCTION

The role of the gut microbiome on host traits has been of interest to many researchers, and it has been connected to issues such as host obesity (Tilg and Kaser, 2011), allergies (McKenzie et al., 2017), and mental health (Du Toit, 2019; Lucas, 2018). Additionally, the importance of gut microbiome in evolutionary biology including role in metabolism, pathogen susceptibility and adaptation has been discussed (Alberdi et al., 2016; Hird, 2017; Kopac and Klassen, 2016) and the biological mechanisms of host-microbiome interactions have been debated (Rosenberg and Zilber-Rosenberg, 2018; Zilber-Rosenberg and Rosenberg, 2008). However, majority of the studies are relying on captive-bred species, while understanding the role of the gut microbiome in eco-evolutionary research requires studying associations between host microbiome and environmental variation in natural environmental conditions, across large scales among and within populations and across taxa.

Interestingly, previous studies have found that there is large-scale intraspecific variation in gut microbiome across populations (Rothschild et al., 2018; Sullam et al., 2012). Population-level differences in gut microbiome have been demonstrated in various taxa, including humans (Gilbert et al., 2018), wild red squirrels (Ren et al., 2017), brown frogs (Tong et al., 2020), and several insect (Sabree and Moran, 2014) and fish species (Liu et al., 2016; Sullam et al., 2015, 2012). However, the environmental drivers behind the population differences are not always well understood. Furthermore, whereas mammalian gut microbiomes are largely defined by phylogeny, many studies have highlighted that environmental variation is likely more important for explaining gut microbiome variation in other taxa, especially birds (Loo et al., 2019).

Birds have been largely neglected in microbiome research and only the recent years have shown an increasing interest in gut microbiome studies with (wild) birds (Bodawatta et al., 2022a; Waite and Taylor, 2014). Birds are a good model species for gut microbiome studies because 1) birds inhabit every continent on Earth and their diversity is well documented (Bibby, 1999; Pereira and Cooper, 2006; Pigot et al., 2020; Rahbek and Graves, 2001). As a result of bird species’ dispersal across the Earth and the biannual migration for some species, birds have developed ways to adapt to a wide range of environmental conditions (Gregory et al., 2005; Koskimies, 1989). This makes them an interesting taxon for studying different mechanism, such as patterns in the gut microbiome, associated with environmental variation (Grond et al., 2018). 2) Within a species, populations are known to differ in phenotype (Charmantier et al., 2008; Husby et al., 2010), and the gut microbiome may contribute to this phenotypic variation among populations. 3) Due to life-history traits such as egg laying, powered flight and migration, the avian gut microbiome may be different from that of e.g., mammals (Grond et al., 2018). Distinct morphological characteristics and the ability to fly have resulted in a high-energy requirement and fast metabolism both of which are influenced by the gut microbiome (Kohl, 2012). Yet, surprisingly, large-scale studies focusing on among-population variation, and the environmental variables explaining variation in the gut microbiome of wild birds among and within populations are still poorly studied (Capunitan et al., 2020; Hird et al., 2015).

Population level differences in avian gut microbiomes could be a result of environmental factors such as season (Davenport et al., 2014; Góngora et al., 2021), diet (Singh et al., 2017), habitat (Drobniak et al., 2022; Loo et al., 2019), temperature (Sepulveda and Moeller, 2020), and humidity (Tajima et al., 2007). Of these factors, season is tightly connected to the other factors and diet is often correlated with habitat. For example, there is a strong seasonal change in the gut microbiome composition of wild mice, which has been suspected to be a result of the transition from an insect to a seed-based diet (Maurice et al., 2015). In thick-billed murres *Uria lomvia* variation in the gut microbiome across the breeding season was explained by prey specialization and differences in diet and sex during the breeding season (Góngora et al., 2021). Similar effect was found in barn swallows *Hirundo rustica*: The swallow diet varied across the breeding season and was correlated with gut microbiome (Schmiedová et al., 2022). In birds, the associations between habitat characteristics and gut microbiome have been studied to some extent: For example, in blue tits *Cyanistes caeruleus* a population living in dense deciduous forests had a higher gut microbiome diversity than a population inhabiting open areas and hay meadows. This may be explained by dense forests having higher overall species abundance and therefore, food item diversity (i.e., diet) and abundance (Drobniak et al., 2022). Diet is also shown to have a positive effect on eastern bluebirds’ *Sialia sialis* nestling gut microbiome: Food supplementation increased the relative abundance of *Clostridium* spp. and was positively correlated with antibody response and lower parasite abundance, thus increasing nestling survival (Knutie, 2020).

Among the abiotic environmental factors, the association between temperature and humidity and the gut microbiome have also been studied, but mostly in other taxa than birds. This study focuses on endothermic species (for ectothermic species see e.g., Bestion et al., 2017; Fontaine et al., 2018; Kohl and Yahn, 2016; Moeller et al., 2020), which maintain their body temperature by generating heat via metabolism (Chevalier et al., 2015; Rosenberg and Zilber-Rosenberg, 2016). Part of this temperature maintenance has been connected to the gut microbiome: The gut microbiome composition of cold exposed laboratory-bred mice *Mus musculus* changed to so called cold microbiota, potentially helping the host to tolerate periods of higher energy demand (Chevalier et al., 2015). In another study with laboratory-bred mice a change in temperature and humidity together with the exposure to wild environment led to different gut microbiome composition than their wild and lab-bred counterparts that resided in lower temperature and humidity (Bär et al., 2020). These changes in the gut microbiome can mediate changes at molecular level and thus, enable adaptation to varying environmental conditions. Temperature has also been shown to have effects on poultry gut microbiomes. Higher temperature can lead to increased gut microbiome species richness and significantly different gut microbiome composition (Wang et al., 2018), and lower temperatures correlate with changes in bacterial composition and muscle amino acid deposition (Yang et al., 2021). In domestic Shaoxing ducks *Anas platyrhynchos* exposure to higher temperatures increased gut microbial abundance and changed the metabolic and transcription-related pathways, which suggests that gut microbiome may have enabled host adaptation to a new thermal environment (Tian et al., 2020). Recent work with wild birds has also shown associations between temperature, host gut microbiome, and host health, although the results are not conclusive (Dietz et al., 2022; Ingala et al., 2021).

The overall aim of this explorative study was to characterize variation in the gut microbiome of wild adult great tit *Parus major* populations across the species’ distribution range in Europe. The great tit is a well-known study species in the fields of ecology and evolution (Krebs, 1971) and provides an attractive study system as this species inhabits vast geographical areas and lives in highly seasonal environments, thus offering the possibility to study the drivers that affect seasonal and population level variation in the gut microbiome. Here, we investigated how 1) population and season contribute to the gut microbiome, and 2) how environmental factors associated with population and season, such as latitude, habitat, rainfall, temperature, and winter diet contribute to the gut microbiome. We expected to see larger seasonal differences in populations living at higher latitudes because abiotic environmental conditions such as snow coverage, rainfall, and temperature vary more towards the polar regions (Anderson and Jetz, 2005; Williams et al., 2015). We predicted that summer season would result in higher gut microbiome diversity, because food abundance, diversity, and time for food foraging is generally higher during summer than winter (Cody, 1981; Karr, 1976). Connecting to this, we also predicted that environmental factors that may be population-specific such as habitat, temperature and humidity significantly contribute to the gut microbiome (Lewis et al., 2017; Murray et al., 2020). For example, a more biodiverse habitat may offer more diverse and abundant prey items and warmer temperatures and moderate rainfall a higher insect abundance, which could lead to higher microbiome diversity and differences in composition (Cox et al., 2019). To our knowledge this is the first study to characterize how environmental variation at a biogeographical scale shapes the variation of the gut microbiome in a wild bird.

## METHODS

### Study area

Fecal samples were collected from wild adult great tits across Europe during winter (January and February) and summer (May and June, breeding season) in 2021 from eight different locations (Fig. 1, SI 1). The aim was to collect samples from ca 20-25 individuals from winter and 20-25 from summer from each location. Due to difficult winter conditions such as colder temperatures and deeper snow coverage, we failed in collecting winter samples from the Westerheide and La Hiruela populations. In total we collected 285 samples, of which 124 samples from winter and 161 samples from summer.

**Figure 1.**
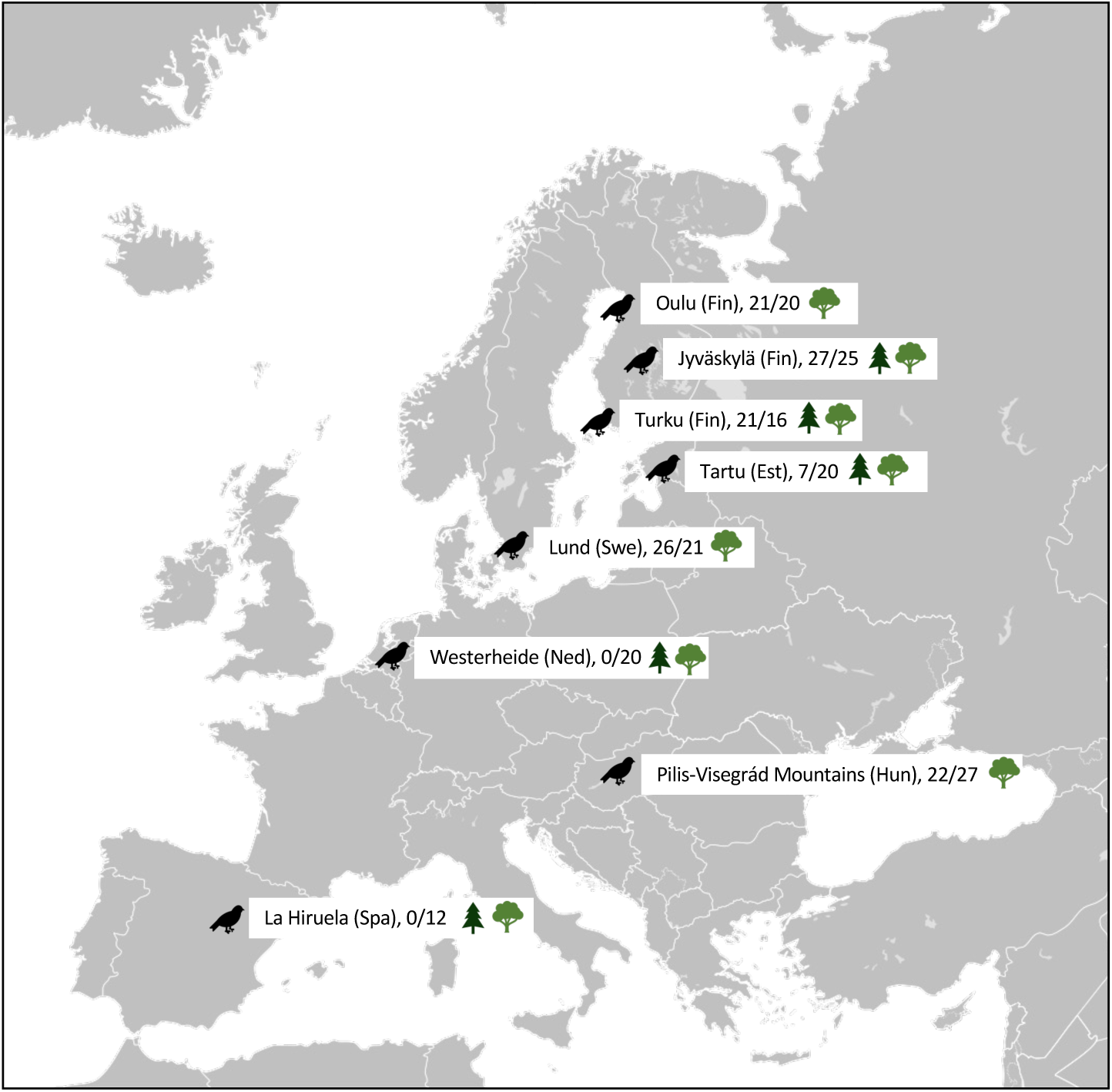
Locations, sample sizes (winter and summer) and habitat types of the eight different great tit populations across the species’ distribution range.

### Fecal sample collection

To capture wild great tits, we used mist nets and feeding traps during winter, and nest box traps during summer. Sample collection followed a protocol by Knutie and Gotanda (Knutie and Gotanda, 2018): adult great tits were captured and put inside a paper bag until defecation, which usually took between 5 and 15 minutes. Fecal samples were then placed straight into 1.5 ml Eppendorf tubes and kept on ice until they were placed in long-term storage in a −80 °C freezer. Each bird was also ringed for identification, sexed, weighed (~0.1g), and their wing-length was measured with a metal ruler (~1mm). Habitat characteristics and latitude were recorded at each population, and temperature (average ambient temperature) and rainfall (average mm per day) data from 2 weeks prior to sampling of each individual bird were collected from the European Climate Assessment and Dataset (https://www.ecad.eu/dailydata/index.php), using the nearest weather station to the sampling location. Winter diet was recorded by monitoring the supplied food at the feeding stations, where birds were caught (SI 1). We did not record summer diet because we could not accurately monitor dietary intake and due to cost limitations. Permits for capturing birds and sample collection were acquired by collaborators at each population.

### DNA extraction and sequencing

We extracted DNA from the collected fecal samples using the Qiagen PowerFecal Pro Kit and followed the manufacturers protocol with minor adjustments: We added a 10-minute incubation step at 65 °C prior to lysis step and used a double elution (eluent was put through the filter twice) to improve DNA yield. To control for contamination and bias during DNA extraction, we included one negative control to each extraction batch and distributed samples from different populations to each extraction batch equally. After extraction, the V4 region of the 16S rRNA gene (approx. length 254 bp) was amplified using the following primers: 515F_Parada (5’-GTGYCAGCMGCCGCGGTAA-3′) (Parada et al., 2016) and 806R_Apprill (5′-GGACT ACNVGGGTWTCTAAT–3′) (Apprill et al., 2015). A total volume of 12 microliters was used in PCR reactions with MyTaq RedMix DNA polymerase (Meridian Bioscience; Cincinnati, OH, USA). We used the following PCR protocol: 1) an initial denaturation at 95 °C for 3 minutes 2) 30 cycles of 95 °C for 45 sec., 55 °C for 60 sec., and 72 °C for 90 sec., and 3) a 10-minute extension at 72 °C at the end. After the first round of PCR, a second round was conducted to apply barcodes for sample identification. For this the protocol was 1) initial denaturation at 95 °C for 3 minutes, 2) 18 cycles of 98 °C for 20 sec., 60 °C for 15 sec., and 72 °C for 30 sec., and 3) final extension at 72 °C for 3 minutes. Each PCR plate also contained a negative control to control for contamination and a ZymoBIOMICS community standard (Zymo Research Corp, Irvine CA, USA) to ensure successful amplification. PCR products’ DNA concentration was measured with Quant-IT PicoGreen dsDNA Assay Kit (ThermoFischer Scientific; Waltham, MA, USA) and quality was checked with gel electrophoresis (1.5 % TAE agarose gel). PCR products were then pooled equimolarly and purified using NucleoMag NGS Clean-up and Size Select beads (Macherey-Nagel; Düren, Germany). Finally, the pools were sequenced with Illumina Novaseq 6000 2 x 250 bp (San Diego, CA, USA) at the Finnish Functional Genomic Center at the University of Turku (Turku, Finland).

### Bioinformatics

#### Sequence processing

The demultiplexed sequence data was processed with QIIME2 version 2021.11 (Bolyen et al., 2018) following the 16S rRNA gene V4 region sequence processing protocol. Adapters were removed using the Cutadapt plugin version 4.4 (Martin, 2011) and quality scores were visually inspected. We used the DADA2 plugin version 2021.4.0 (Callahan et al., 2016) to truncate reads at 220 bp and to generate amplicon sequence variants (hereafter ASVs), which stand for each individual bacterial sequence (Eren et al., 2013). We used the SILVA v132 database with the sk-learn classifier to assign taxonomy (Quast et al., 2013; Yilmaz et al., 2014). We used the phylogeny plugin to construct a rooted phylogenetic tree, and removed singletons, eukaryotes, mitochondria, archaea, chloroplasts, and unassigned taxa in QIIME2 before further analysis. We then combined the resulting ASV table with metadata, taxonomy table and phylogenetic tree using the *phyloseq* package version 1.44.0 (McMurdie and Holmes, 2013) in R program version 4.3.0 (R Core Team). Contaminants (N=61 ASVs) were removed using the *decontam* package version 1.20.0 (Davis et al., 2018). We also filtered samples that had less than 100 reads as they were likely a result of an error in amplification. The resulting dataset had 15 288 ASVs in 284 samples (total number of reads in the whole dataset 16 629 323, average number of reads per sample 57 740, median number of reads 17 189) (SI 2).

For downstream analyses of gut microbiome diversity (i.e., alpha diversity), the dataset was rarified at 1000 reads based on the level at which the rarefaction curves plateaued. This was conducted to account for uneven sequencing depth between samples to normalize the data and to avoid the bias that rare taxa may have in the analyses (Cameron et al., 2021; Schloss, 2023; Weinroth et al., 2022). Seven samples were excluded from the dataset in rarefying resulting in a total of 277 samples and 6883 ASVs, which divided into 121 winter samples and 156 summer samples. We tested both the rarefied and unrarefied datasets for consistency in gut microbiome diversity. For analyses of gut microbiome composition (i.e., beta diversity), we used the unrarefied dataset. For both gut microbiome diversity and composition analyses, we checked that the results were consistent between the unrarefied and rarefied datasets.

### Data analysis

#### Gut microbiome diversity

We used Shannon Diversity Index and Chao1 Richness (Chao, 2006) as the gut microbiome diversity (i.e., alpha diversity) metrics using the *phyloseq* package version 1.44.0 (McMurdie and Holmes, 2013). In each model we first ran the model with Shannon Diversity Index as the response variable and then with Chao1 Richness. We use these two metrics because Shannon Diversity Index considers both taxa abundance and evenness and Chao1 Richness measures the observed number of taxa. Chao1 Richness is more sensitive to rare taxa, whereas Shannon Diversity Index is more robust as it is not easily affected by the presence of rare taxa (Haegeman et al., 2013). For all gut microbiome diversity analyses, we use the rarefied dataset (N=277). Additionally, we use both body condition (linear regression residual of weigh ~ wing) and weight as proxies for individual condition: we ran each model first with body condition and then with weight replacing body condition. Because some birds escaped prior to measurements, and in one population wing length was not recorded, we do not have a weight and wing measurement for every bird in this study (total of 46 birds from 3 different populations).

All statistical analyses were conducted in R program version 4.3.0 (R Core Team). Normality and homoscedasticity of the residuals were visually assessed. Variance Inflation Factors were assessed for each model with the package *DHARMa* version 0.4.6 (Hartig and Hartig 2017). Linear models were conducted using the packages *lme4* version 1.1-33 (Bates et al. 2014) and *car* version 3.1.2 (Fox et al. 2012).

First, we used a linear model to test if season (categories: winter and summer) and population (6 categories) contribute to the gut microbiome diversity in all the observed populations. We used gut microbiome diversity as the response variable and population and season as the predicting variables. We also ran this same model with an interaction between season and population to test for population differences across seasons (SI 5). In these models, the Westerheide and La Hiruela populations were excluded as those populations were only measured during summer and including them may bias the results. Oulu was set as the population reference level because it was the northernmost of our study populations. None of the explanatory factors were correlated (VIFs<4).

Second, we tested in more detail, which environmental factors across and within populations and seasons associate with the variation in gut microbiome diversity. We ran a linear model with gut microbiome diversity as the response variable and the following fixed factors: latitude (continuous variable), habitat (categories: mixed and deciduous), rainfall (continuous variable), and temperature (continuous variable) using data across both seasons. Sex (category variable) and body condition / weight (continuous variable) were also included in the model as fixed factors to control for individual differences within population because physiological factors may contribute to variation in the gut microbiome (Amato et al., 2019; Corl et al., 2020; Góngora et al., 2021; Jašarević et al., 2016; Ley et al., 2008; Zhao et al., 2013). In these models, we excluded the Westerheide and La Hiruela populations as those populations were only recorded during summer. None of the explanatory factors were correlated (VIFs<4).

Third, because of the uneven sample sizes for winter and summer observations and because diet was only monitored during winter, we analyzed gut microbiome diversity separately by season. For winter data (N=121), we ran a linear model to analyze whether latitude, habitat, temperature, rainfall, and diet (categories: sunflower seeds and sunflower seeds + peanuts) contribute to gut microbiome diversity in populations during winter. Again, body condition / weight and sex were also included in the model. The winter model included diet as an explanatory variable as we had recording of winter feeding. None of the explanatory factors were correlated (VIFs<4).

For summer data (N=156) we ran a similar model as we did for the winter data. We used gut microbiome diversity as the response variable and latitude, habitat, rainfall, temperature, and body condition / weight and sex as explanatory variables. We did not include diet in the model because we did not record summer diet. None of the explanatory factors were correlated (VIF<4).

For each model, we tested the significance factors using *F*-test ratios in analysis of variance (ANOVA type 3). Ideally, non-independence of individuals within a population should be controlled for by using linear mixed effects models and population as a random effect. Yet, when such models were performed (packages *lme4*, version 1.1-33, Bates et al. 2014; *car*, version 3.1.2, Fox et al. 2012), the model did not converge, as many of the environmental factors were measured at the population level.

#### Gut microbiome composition

For gut microbiome composition (i.e., beta diversity), we used the *microbiome* package version 1.22.0 (Lahti and Shetty 2017). We visualized the gut microbiome compositions between populations and seasons with non-metric multidimensional scaling (NMDS). For these visualizations, we used the Bray-Curtis dissimilarity metric that examines the dissimilarity of microbes among samples (Bray and Curtis, 1957). To analyze variation in gut microbiome communities among populations, we used permutational analysis of variance (PERMANOVA; *vegan* package, version 2.6-4, Oksanen et al. 2013) with the *adonis2* function and 9999 permutations. We constructed these PERMANOVA models the same way as we did the multiple linear regression models for the gut microbiome diversity measurements. First, we analyzed whether season and population contribute to the variation in gut microbiome composition. Second, we analyzed whether latitude, body condition / weight, habitat, rainfall and temperature, and sex contribute to the variation in gut microbiome composition across both seasons. Third, we used the winter and summer data subsets to analyze associations between the gut microbiome composition and environmental variables within season. We tested the homogeneity of variance (beta dispersion), which showed similar dispersion for populations (BETADISPER^9999^, F_5_=1.104, P=0.374) and seasons (BETADISPER^9999^, F_1_=1.417, P=0.235), thus affirming that PERMANOVA is appropriate for comparing community compositions.

We also ran a differential abundance analysis (DESeq2) to see whether there are differences in bacterial taxa abundance within populations between winter and summer. For this analysis, we only used the populations that have both winter and summer data (N=6 populations) because the aim of this analysis is to compare within population differences in taxa abundance between seasons. All taxa are identified at least to family level and some to genus level. Unfortunately, many of these observed taxa are less studied and their functions in the gastrointestinal tract, especially beyond humans, are not known. Here we focus on the taxa that are more studied in gut microbiome research. For visualizing the DESeq2 results, we used order level to make the plot readable. We used the package *DESeq2* version 1.40.1 (Love et al., 2014) for the differential abundance analysis.

## RESULTS

### Gut microbiome diversity among populations and across both seasons

There were 27 bacterial phyla detected across all samples, and the most abundant phyla were *Proteobacteria*, *Actinobacteria*, and *Firmicutes*. While there was variation in gut microbiome between populations, population did not significantly influence gut microbiome diversity (Fig. 2A, Table 1, SI 4). Season significantly influenced gut microbiome diversity when Shannon Diversity Index was used as the response variable (P=0.011, Fig. 2B, Table 1, SI 4): diversity was higher in winter than in summer. We found no significant interaction between season and population (P all>0.05, SI 5). Furthermore, none of the environmental factors or body condition / weight contributed to gut microbiome diversity when the model included populations from both seasons (Table 1, SI 4). When Chao1 Richness was used as the response variable, none of the explanatory factors significantly contributed to gut microbiome diversity (Table 1, SI 4).

**Figure 2.**
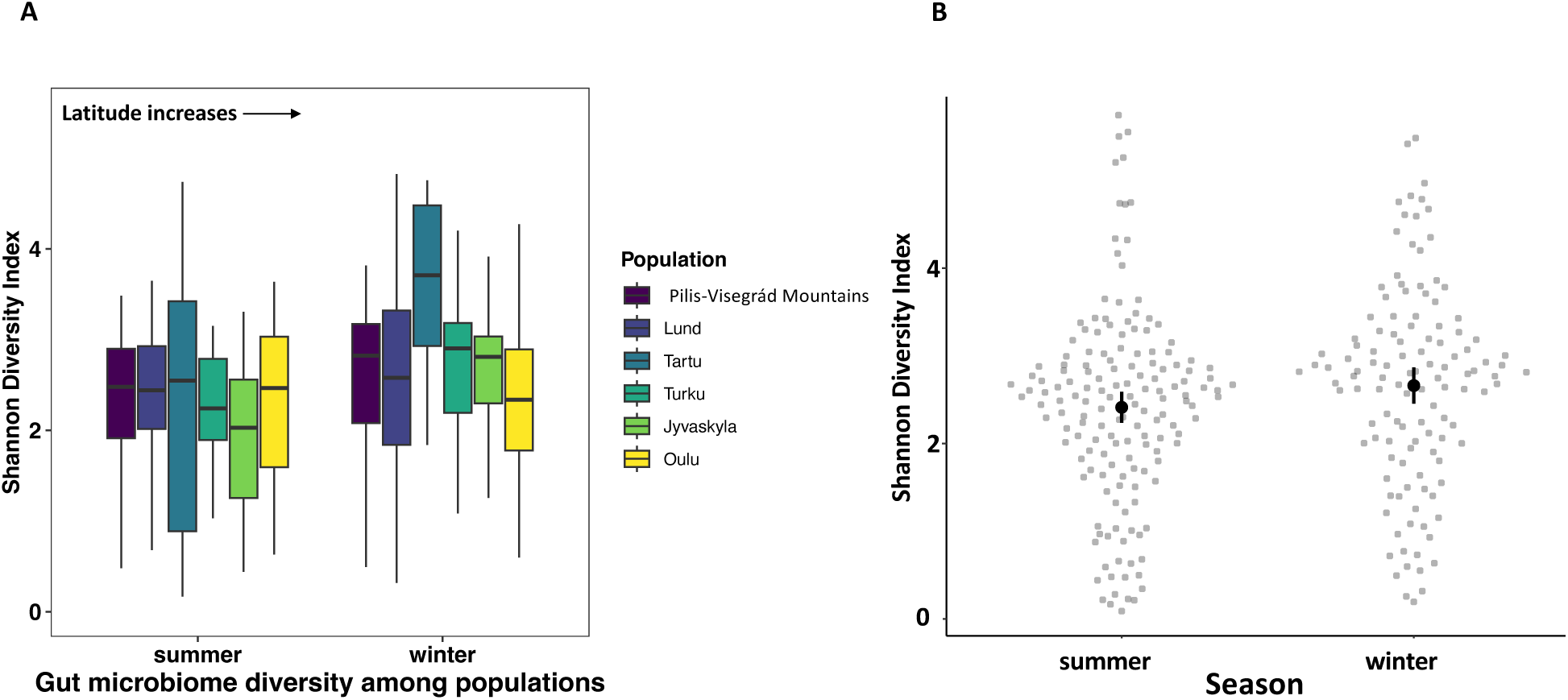
A) Gut microbiome diversity (Shannon Diversity Index) among populations and between seasons ordered by latitude (south to north), and B) the effect of season on gut microbiome diversity (mean and SE)

**Table 1.**
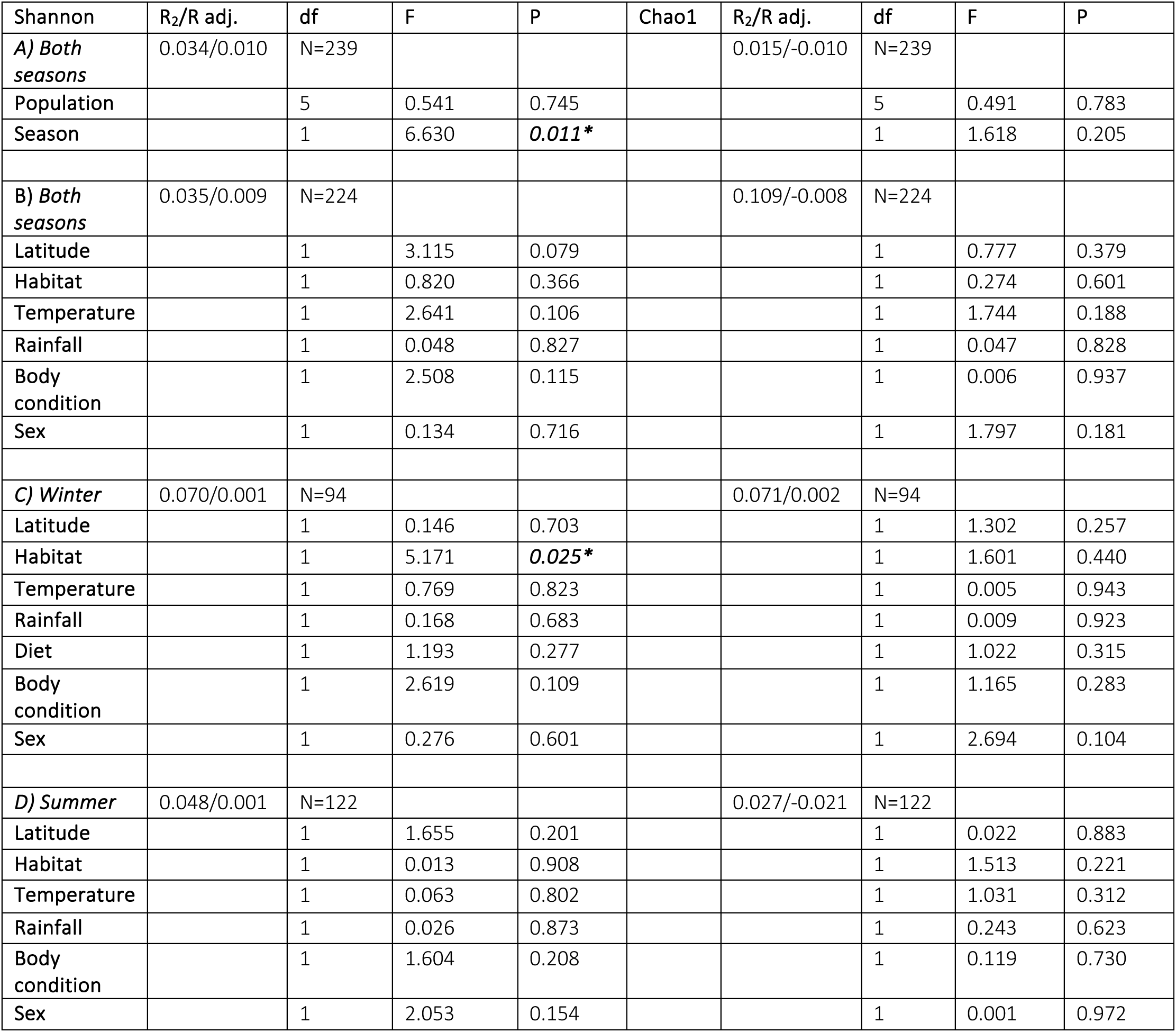
The association between gut microbiome diversity (Shannon and Chao1) and A) population and season, B) latitude, habitat, temperature, rainfall, body condition and sex between seasons, C) latitude, habitat, temperature, rainfall, diet, body condition and sex during winter, and D) latitude, habitat, temperature, rainfall, body condition and sex during summer. Linear model analyses were used and the ANOVA output with Satterthwaite’s method is reported in the table.

### Winter subset

In winter, gut microbiome diversity was higher in individuals inhabiting mixed forests than deciduous forests when measured with Shannon Diversity Index (P=0.025, Fig. 3, Table 1, SI 4), but not when measured with Chao1 Richness (Table 1, SI 4). The result was the same for habitat when body condition was replaced with weight in the model (Shannon P=0.033, SI 9; Chao1 P=0.218, SI 4). Latitude, temperature, rainfall, and diet (P all>0.05) did not contribute to gut microbiome diversity in any of the models (Table 1, see SI 4).

**Figure 3.**
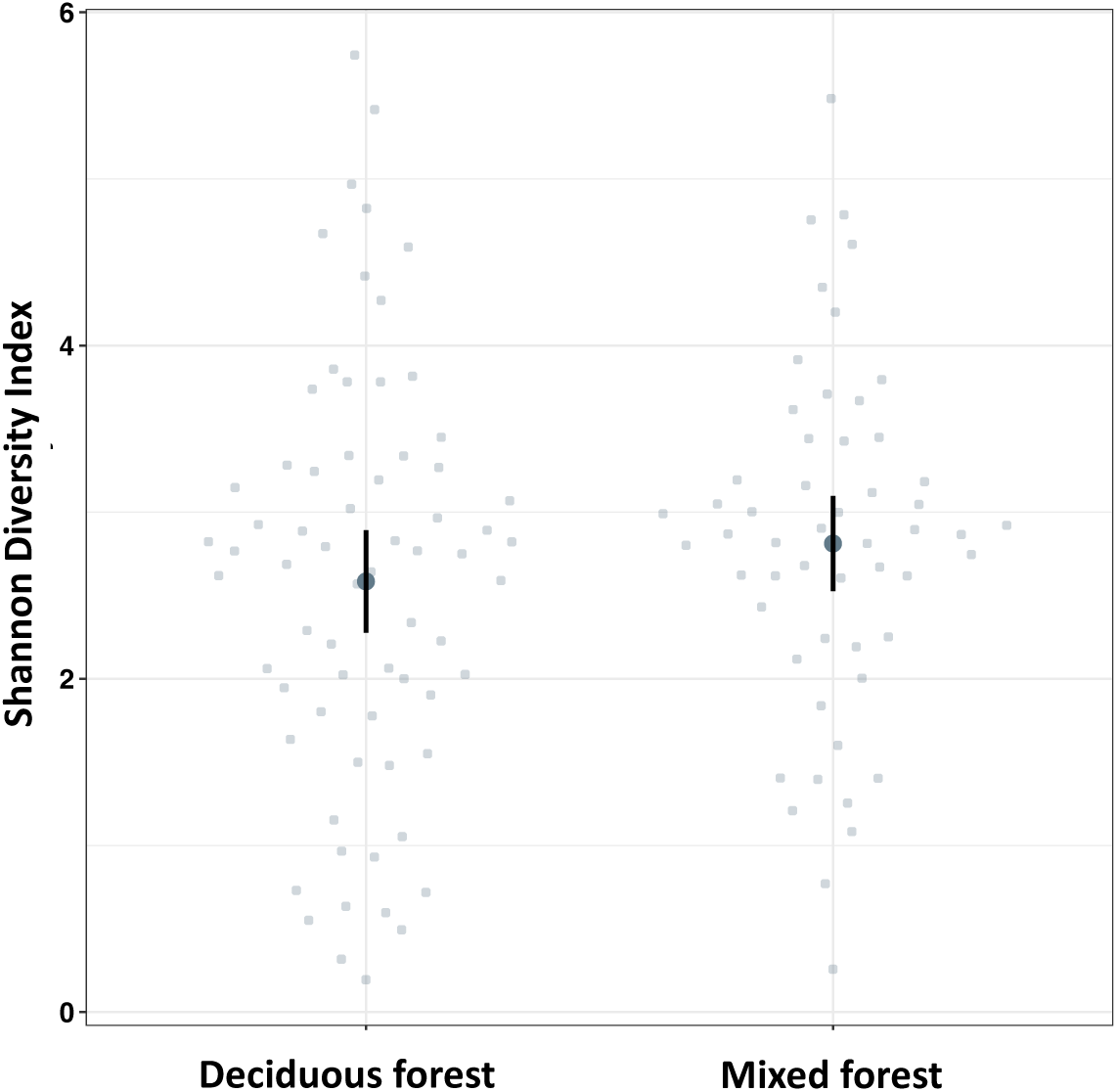
The effect of habitat on gut microbiome diversity (mean and SE) during winter. Populations inhabiting deciduous habitat are Oulu, Lund and Pilis-Visegrád Mountains, and populations inhabiting mixed habitat are Jyväskylä, Turku and Tartu.

### Summer subset

Neither latitude, habitat, temperature, and rainfall nor body condition / weight and sex contributed to gut microbiome diversity during summer (P all>0.05, Table 1, see SI 4).

### Gut microbiome composition

As with gut microbiome diversity, visual observation of the gut microbiome composition showed that there was population level variation in composition (Fig. 4). However, PERMANOVA showed that population did not significantly contribute to differences in gut microbiome composition among populations (R^2^=0.021, P=0.397, Fig. 4, SI 6).

**Figure 4.**
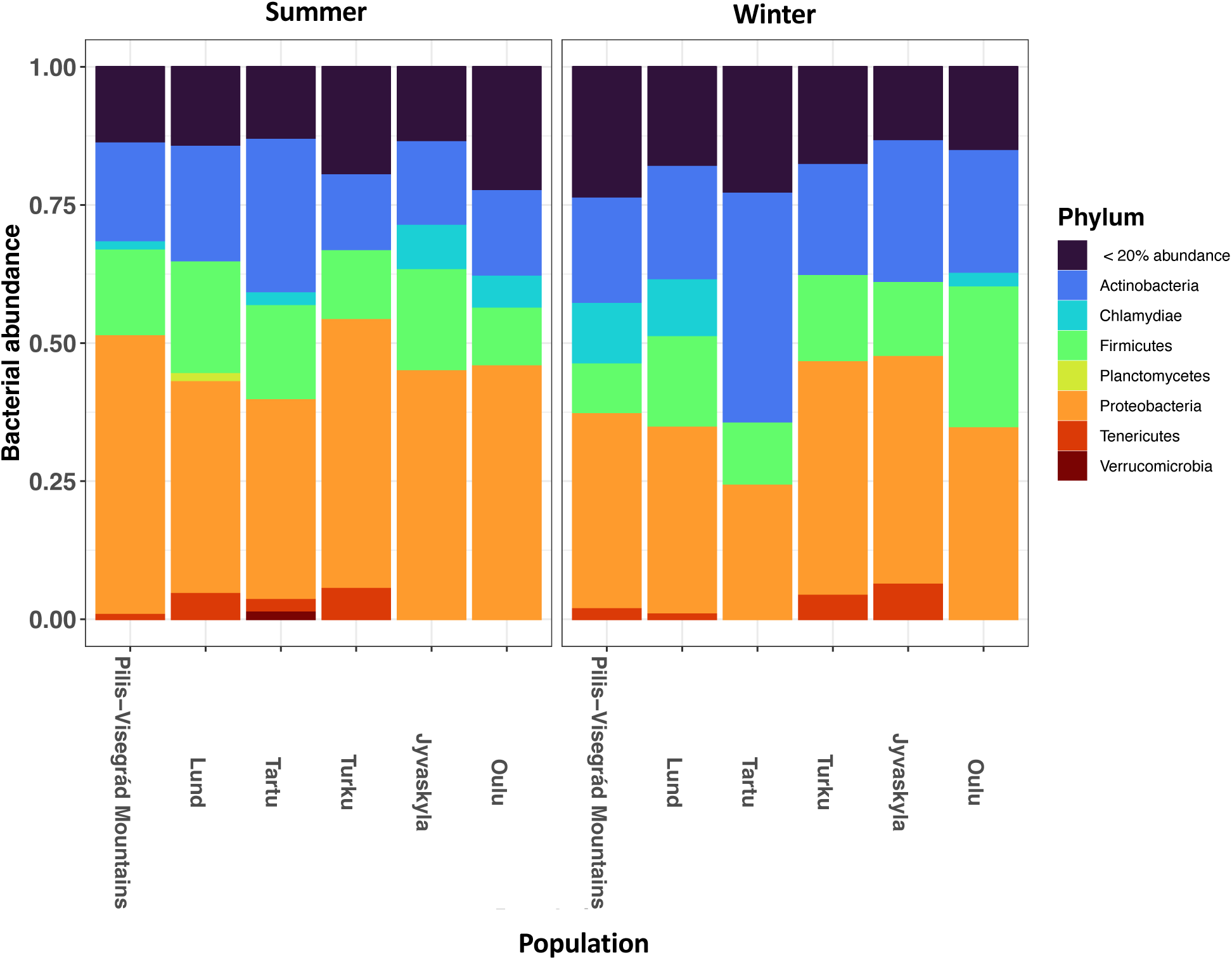
Among-population comparison of gut microbiome relative abundance on phylum level between seasons. Less abundant phyla are summed up as “< 20% abundance” to improve plot readability.

PERMANOVA showed that there were significant differences in composition between seasons, but season only explained 0.5% of these differences (R^2^=0.005, P=0.034, SI 6). Of the environmental factors temperature explained 0.6% of differences in gut microbiome composition in all data across both seasons (R^2^=0.006, P=0.012, Fig. 5, SI 6). When looking at the winter and summer subsets of data, none of the measured factors explained the differences in gut microbiome composition (P all>0.05, SI 6)

**Figure 5.**
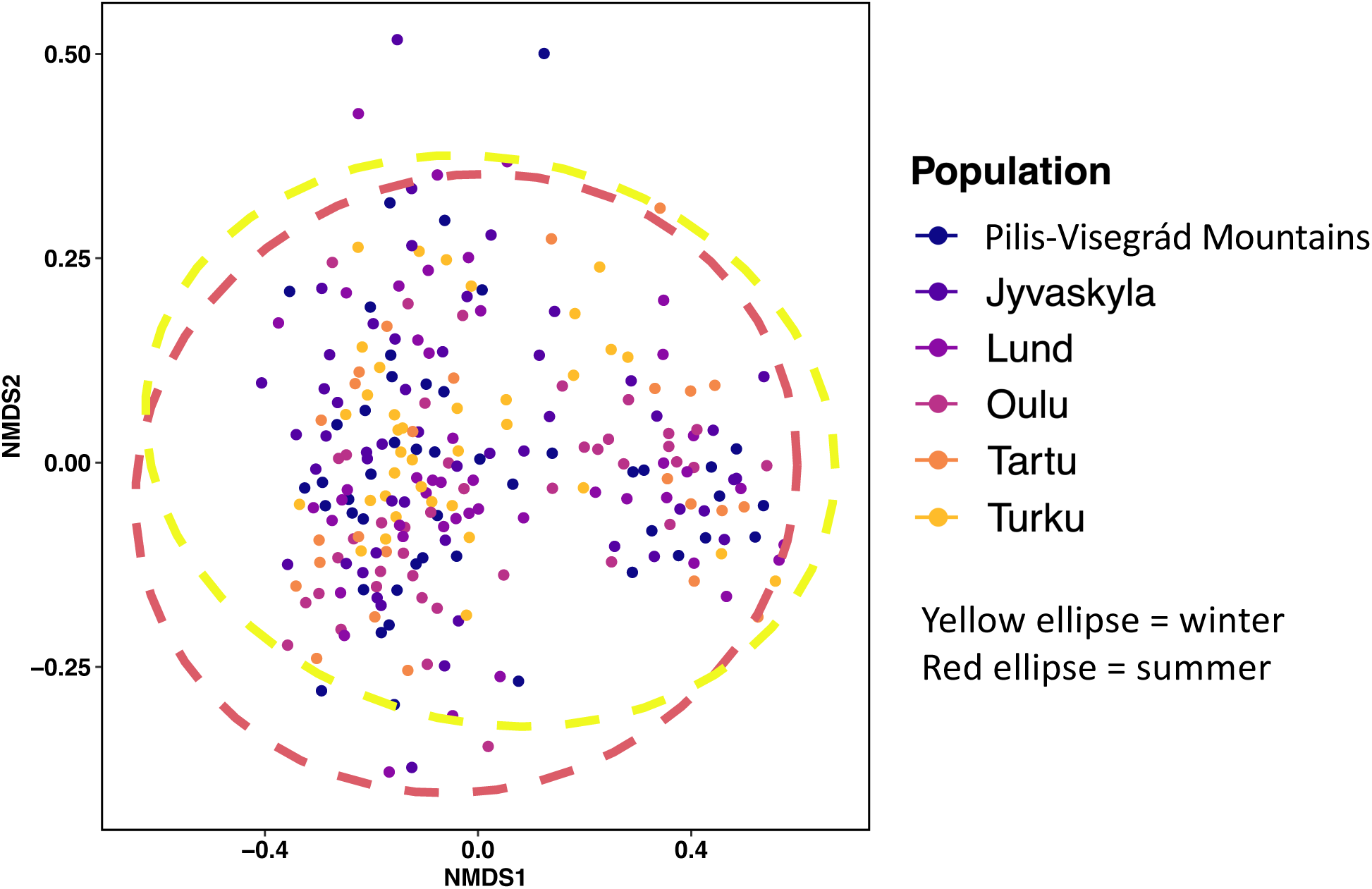
Non-metric multidimensional scaling measured with Bray-Curtis dissimilarity representing the microbial composition dissimilarity among populations between winter and summer. Points are colored according to population. Ellipses represent 95 % confidence intervals.

Seasonal differences in bacterial taxa abundance were detected in each population (Fig. 6, SI 7), and some of these taxa were of interest to us due to their known beneficial or pathogenic effects. Of the well-known taxa, the order *Bacillales* were more abundant in Pilis-Visegrád Mountains, Turku and Jyväskylä during summer than winter, and more abundant in Oulu during winter than summer. The order *Bifidobacteriales* were more abundant in Turku during winter than summer. The order *Chlamydiales* were more abundant in Turku and Jyväskylä during summer than winter. The order *Enterobacteriales* were more abundant in Oulu during summer than winter. The order *Lactobacillales* were more abundant in Pilis-Visegrád Mountains during summer than winter and in Turku during winter than summer. The order *Micrococcales* were more abundant in Tartu, Lund, Pilis-Visegrád Mountains, Oulu, and Turku during winter than summer.

**Figure 6.**
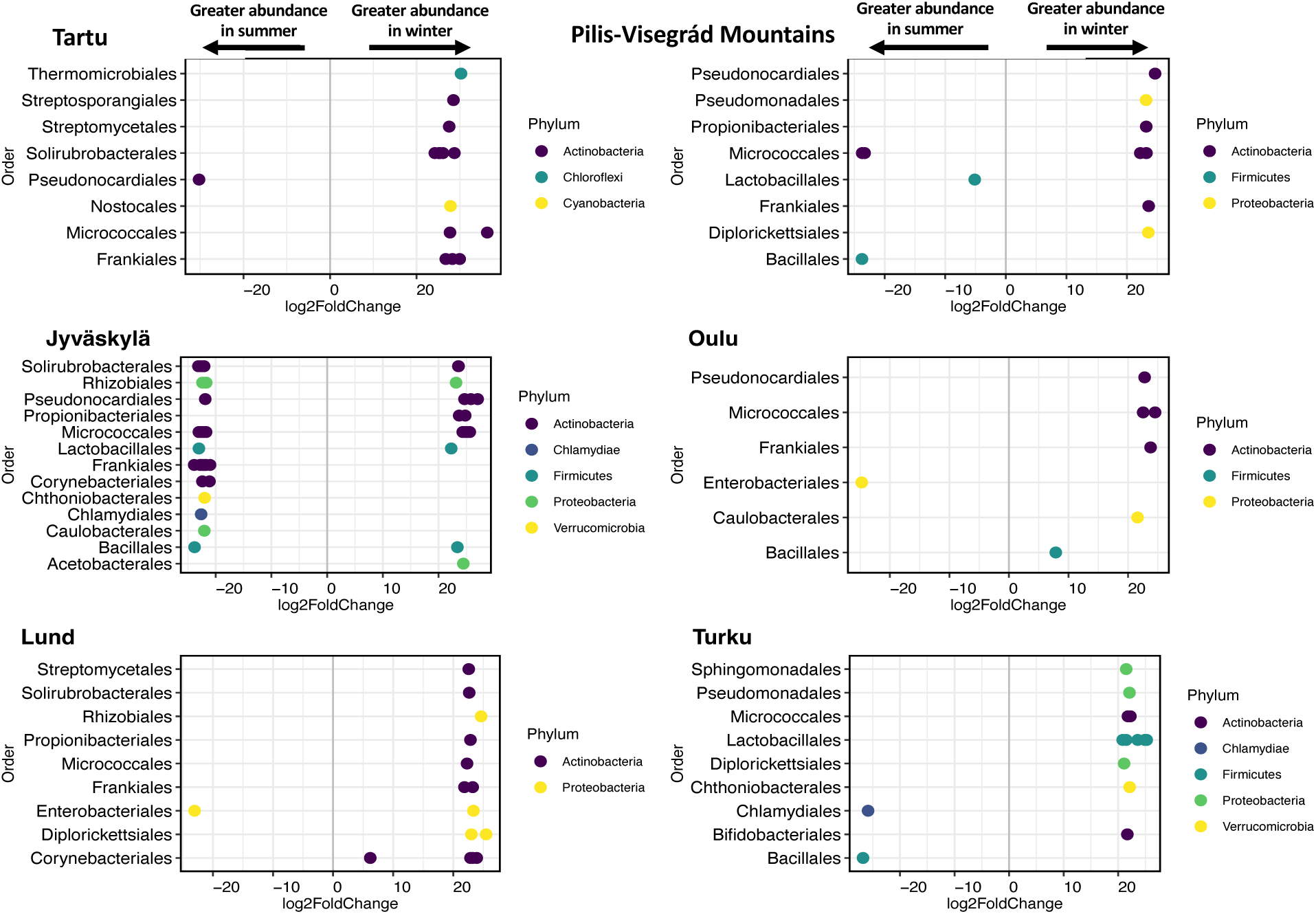
Visualization of the differential abundance analysis comparing six great tit populations between winter (positive log2FoldChange) and summer (negative log2FoldChange). Each dot represents one taxon within a bacterial order. All taxa are identified at least to Family level (SI 7), but for figure readability they are plotted on order level.

## DISCUSSION

The goal of this study was to characterize large-scale variation in wild adult great tit gut microbiomes and analyze whether environmental factors associated with population and season associate with the gut microbiome. The majority of bacterial taxa in our samples belonged to the phyla *Proteobacteria*, *Firmicutes* and *Actinobacteria*, which was expected as they are the key phyla of great tit gut microbiome (Bodawatta et al., 2021a; Teyssier et al., 2018a). We did not find large-scale among population variation in the gut microbiome diversity or composition. Instead, we found the gut microbiome diversity and composition to be dependent on seasons and diversity to be dependent on habitats during the winter season. Importantly, while we did not find evidence for the effects of latitude, weather, or individual characteristics on gut microbiomes, much of the variation was left unexplained suggesting unknown sources of variation.

### No major differences among populations in microbiome diversity or composition

We found no evidence for population differences in the gut microbiome diversity or composition. This is surprising in light of apparent differences in environmental factors and given that great tits also show population-level differences in physiology (Saulnier et al., 2023) phenotypes (Dingemanse et al., 2012; Gamero et al., 2015), and minor genetic differentiation (Lemoine et al., 2016; Noordwijk et al., 2002), which have been shown to contribute to among-population variation in the gut microbiome of various taxa (Gadau et al., 2019; Meng et al., 2014; Spor et al., 2011; S. Wang et al., 2018; Wen et al., 2021). We observed high within-population variation in gut microbiome diversity, which was expected because especially in smaller bird species individual variation in gut microbiome diversity has been found to be very high (as mentioned in Bodawatta et al., 2021b). In our study, all populations were in forested habitats with high plant species diversity, which could explain why large among-population differences in the gut microbiome were not observed (compared to for example the observed microbiome differences between urban and rural habitats (Phillips et al., 2018; Teyssier et al., 2020)).

### Gut microbiome diversity and composition differ across seasons

We found that gut microbiome diversity is higher in great tit populations during winter than summer, and that gut microbiome composition varies between seasons. This is in line with many studies reporting seasonal variation in the gut microbiome (Baniel et al., 2021; Davenport et al., 2014; Góngora et al., 2021; Ren et al., 2017; Xiao et al., 2019). However, we expected that the gut microbiome diversity and composition would be lower during winter due to limited foraging times and the breadth of available dietary items for great tits (Grubb, 1978; McNamara et al., 1994; Vel’ký et al., 2011). During winter, great tits can use both insects (lepidopterans, coleopterans and dipterans), plant material (seeds and buds) and human provided food, compared to only insectivorous diet during summer (Vel’ký et al., 2011). If the birds in our study populations were able to use a high diversity of food items, it would reflect to their gut microbiome and explain why the great tits in our study had higher gut microbiome diversity during winter. The result would also follow some previous studies in which diet diversity has been connected to gut microbiome diversity (Knutie et al., 2019; Teyssier et al., 2018b).

We also observed within-population seasonal shifts in taxa abundances. Of the populations we sampled Oulu, Jyväskylä and Turku were the most northern and experienced the widest range of environmental changes between winter and summer and are therefore expected to show the largest changes. The rest of the populations were located more to the south and west of Europe, which can mean milder seasonal changes in environment and less snow cover (Baker., 1939). Of the six populations compared here, Jyväskylä showed the most differences in between-season taxa abundance and Oulu the least differences, which was opposite to what we expected. The order *Enterobacteriales* was more abundant in Lund, Pilis-Visegrád Mountains, and Oulu during summer than winter. Many bacterial taxa belonging to the order *Enterobacteriales* such as *Salmonella enterica* and *Escherichia coli* are known pathogens in birds (Cheville and Arp, 1978; Tizard, 2004). The order *Chlamydiales* was more abundant in Jyväskylä and Turku during summer than winter. These pathogens are likely to be more abundant during summer, because individual birds can pass them on to other individuals during copulation (Escallón et al., 2019; Grond et al., 2018). The order *Bacillales* (not to be mixed with *Lactobacillales*), which contains several pathogenic genera such as *Staphylococcus*, *Bacillus* and *Listeria*, was also more abundant in Pilis-Visegrád Mountains, Jyväskylä and Turku during summer than winter and more abundant in Oulu during winter than summer. Of the beneficial taxa, the order *Lactobacillales* abundance varied between populations: they were more abundant during winter than summer in Turku and more abundant during summer than winter in Pilis-Visegrád Mountains and Jyväskylä. Especially the genus *Lactobacillus* of the order *Lactobacillales* is known for its importance digestive health (Reid and Burton, 2002), and these beneficial health effects are also known from poultry (Al-Khalaifah, 2018).

### Habitat associates with gut microbiome diversity, but not composition, during winter

Mixed forest associated with higher gut microbiome diversity than deciduous forest during winter, but not during summer. There were no differences in microbiome composition between habitats. Habitats with mixed tree and other plant species promote diversity in forest associated taxa (Ampoorter et al., 2020; Tinya et al., 2021), resulting in a wider range of dietary items for the great tits. A more diverse diet has been found to associate with higher gut microbiome diversity (Bodawatta et al., 2022b) and could also explain why great tits inhabiting mixed forest had more diverse gut microbiomes during winter. Furthermore, breeding greatly influences physiology (Norte et al., 2010) and gut microbiome diversity (Escallón et al., 2019; Góngora et al., 2021; Zheng et al., 2020). Such physiological changes could overrun effects of the environment, such as the habitat, in the samples collected during the breeding season (but see Drobniak et al., 2022). It also leaves us questioning whether the differences in gut microbiome diversity between habitats would appear later during summer. As breeding comes with a great physiological cost (Norte et al., 2010), the gut microbiome may change prior, during and after the breeding season (Escallón et al., 2019).

### Lack of association between abiotic and intrinsic biotic factors on the gut microbiome variation

We found no association between latitude, rainfall, winter diet, and gut microbiome diversity or body condition / weight, sex, and gut microbiome diversity. However, we did find that temperature was weakly linked to gut microbiome composition, but not diversity. We expected that lower temperature would lead to lower gut microbiome diversity, and influence microbiome composition given previous studies with bird gut microbiomes (Dietz et al., 2022; Ingala et al., 2021; Tian et al., 2020; Wang et al., 2018; Yang et al., 2021) and mammal studies (e.g., Worthmann et al., 2017; Zhang et al., 2018). Great tits are an endothermic species, which most likely means that ambient temperatures may not have a major effect in the gut microbiome diversity / composition (Ingala et al., 2021) even though in mice studies effects between temperature and gut microbiome have been found (Chevalier et al., 2015). However, many of the previous studies were conducted with extreme temperatures and in captive conditions. For example, in egg laying hens spells of extreme hot temperatures lead to a decrease in *Firmicutes* abundance, a taxon that is known for its importance in short-chain fatty acid metabolism (Zhu et al., 2019). It is likely that the slight association between temperature and the gut microbiome composition is a result of the populations being in different parts of Europe and thus, they experience a varying range of temperatures throughout the year.

Furthermore, both rainfall and snowfall can affect food item diversity and abundance and reflect on the gut microbiome diversity (Baniel et al., 2021; Schmiedová et al., 2023). For example, rainfall can influence insect abundances during summer, which are significant dietary items for great tits (Schöll et al., 2016). Severe weather can also limit foraging time leading to temporary depletion in food intake (Brittingham and Temple, 1988). This can result in increased physiological stress that has been shown to impact the gut microbiome diversity (Noguera et al., 2018). However, this limited foraging time may be more reflected on the nestlings (Radford et al., 2001) as the gut microbiome is established at the nestling stage (Davidson et al., 2021; Teyssier et al., 2018a).

We found no association between supplemented winter diet and the gut microbiome. We provided sunflower seeds or peanuts or the mix of those two, which may not significantly change the gut microbiome, and great tits will additionally use a wide variety of other food items across all populations. Finding associations between diet and the gut microbiome requires more fine-tuned experiments such as captive experiments in which dietary items and food intake are carefully monitored. Also, sampling at multiple timepoints could be used to see possible longitudinal changes the gut microbiome (as suggested in Davidson et al., 2021).

Our results concerning body condition / weight are in line with recent studies that have not found a single conclusion between the gut microbiome diversity / composition and body condition. In nestling great tits, one study found that better body condition connected to higher gut microbiome diversity (Teyssier et al., 2018a), whereas in another study there was no association between the two factors (Liukkonen et al., 2023). In adult birds, there was no association between body condition and the gut microbiome diversity in Seychelles’ warblers *Acrocephalus sechellensis* (Worsley et al., 2021) or in white-crowned sparrows *Zonotricia leucophrys* (Phillips et al., 2018). Contrastingly, in adult female steppe buzzards *Buteo buteo vulpinus* body condition associated with higher gut microbiome diversity, but no effect was found in male birds (Thie et al., 2022). In our study, it could be that the birds had generally good body condition as they were the ones that had survived to the first winter post-fledging or to breeding season (Naef-Daenzer et al., 2001; Naef-Daenzer and Grüebler, 2008). It may be beneficial to sample birds at multiple timepoints throughout the year to detect possible longitudinal changes in gut microbiome and body condition. Furthermore, the association between sex and gut microbiome diversity has proven to be inconclusive in previous avian gut microbiome studies. In blue tits sex did not associate with gut microbiome diversity or composition (Drobniak et al., 2022) and similar result was found in barn swallows (Kreisinger et al., 2015). During the breeding season bird species that have multiple sexual partners pass cloacal microbiota during copulation, which could result in more similar gut microbiome samples between sexes (Grond et al., 2018). Also, sex-based differences in bird gut microbiomes may be difficult to detect with restricted sample sizes (Capunitan et al., 2020).

## CONCLUSIONS

This study is among the first to disentangle the large-scale variation in the gut microbiome of wild adult great tits. It provides new results about the dynamics of wild great tit gut microbiomes and how environmental factors such as season, habitat and temperature influence the gut microbiome. Our key finding is that season significantly associates with both gut microbiome diversity and composition and factors such as habitat and temperature, which are largely influenced by season, also associate with the gut microbiome. Our results show that changes in environmental conditions can alter the gut microbiome, thus highlighting the importance of studying the effects of environmental change on gut microbiomes. More work is needed to understand the origins of the observed within and among-population variation in great tit gut microbiomes and how this variation connects to population performance in changing environmental conditions.

## AUTHOR CONTRIBUTION

Martta Liukkonen planned the project, organized data collection processed the samples, prepared the sequence libraries, analyzed the data, and wrote the manuscript. Suvi Ruuskanen planned and funded the project, helped in sample processing, data analysis and manuscript writing. Kirsten Grond assisted in sequence library preparation protocols, bioinformatics, and data analysis. Jaime Muriel, Jesús Martínez-Padilla, Andreas Nord, Veli-Matti Pakanen, Balázs Rosivall, Kees van Oers collected fecal samples and commented on the manuscript. All authors have approved the final manuscript.

## ACKNOWLEDGEMENTS

For their help in the field, we would like to thank Cassandre Deparde, Juho Jolkkonen, Nelli Leskisenoja and Inka Ojanen for collecting samples in Jyväskylä. Jorma Nurmi (University of Turku) and Antoine Stier (IPHC Strasbourg, University of Turku) for collecting samples on Ruissalo island and Peter de Vries (NIOO-KNAW) for help in collecting samples in Westerheide. We would like to thank Emil Aaltonen Foundation (grant to Suvi Ruuskanen) for funding this research. Andreas Nord was funded by the Swedish Research Council (grant no. 2020-04686).

## DATA AVAILABILITY

Sequence data will be available at NCBI Sequence Read Archive (PRJNA1036439). QIIME2 script and R codes are available at GitHub (https://github.com/marttal/GTvariation) and on request.

## REFERENCES

Alberdi, A., Aizpurua, O., Bohmann, K., Zepeda-Mendoza, M.L., Gilbert, M.T.P., (2016). Do Vertebrate Gut Metagenomes Confer Rapid Ecological Adaptation? Trends in Ecology & Evolution 31, 689–699. 10.1016/j.tree.2016.06.008

Al-Khalaifah, H.S., (2018). Benefits of probiotics and/or prebiotics for antibiotic-reduced poultry. Poultry Science 97, 3807–3815. 10.3382/ps/pey160

Amato, K.R., G. Sanders, J., Song, S.J., Nute, M., Metcalf, J.L., Thompson, L.R., Morton, J.T., Amir, A., J. McKenzie, V., Humphrey, G., Gogul, G., Gaffney, J., L. Baden, A., A. O. Britton, G., P. Cuozzo, F., Di Fiore, A., J. Dominy, N., L. Goldberg, T., Gomez, A., Kowalewski, M.M., J. Lewis, R., Link, A., L. Sauther, M., Tecot, S., A. White, B., E. Nelson, K., M. Stumpf, R., Knight, R., R. Leigh, S., (2019). Evolutionary trends in host physiology outweigh dietary niche in structuring primate gut microbiomes. ISME Journal 13, 576–587. 10.1038/s41396-018-0175-0

Ampoorter, E., Barbaro, L., Jactel, H., Baeten, L., Boberg, J., Carnol, M., Castagneyrol, B., Charbonnier, Y., Dawud, S.M., Deconchat, M., Smedt, P.D., Wandeler, H.D., Guyot, V., Hättenschwiler, S., Joly, F., Koricheva, J., Milligan, H., Muys, B., Nguyen, D., Ratcliffe, S., Raulund-Rasmussen, K., Scherer-Lorenzen, M., Van Der Plas, F., Keer, J.V., Verheyen, K., Vesterdal, L., Allan, E., (2020). Tree diversity is key for promoting the diversity and abundance of forest-associated taxa in Europe. Oikos 129, 133–146. 10.1111/oik.06290

Anderson, K.J., Jetz, W., 2005. The broad-scale ecology of energy expenditure of endotherms. Ecology Letters 8, 310–318. 10.1111/j.1461-0248.2005.00723.x

Apprill, A., McNally, S., Parsons, R., Weber, L., (2015). Minor revision to V4 region SSU rRNA 806R gene primer greatly increases detection of SAR11 bacterioplankton. Aquatic Microbial Ecology 75, 129–137. 10.3354/ame01753

Baker., J.R., 1939. The Relation between Latitude and Breeding Seasons in Birds. Proceedings of the Zoological Society of London A108, 557–582. 10.1111/j.1096-3642.1939.tb00042.x

Baniel, A., Amato, K.R., Beehner, J.C., Bergman, T.J., Mercer, A., Perlman, R.F., Petrullo, L., Reitsema, L., Sams, S., Lu, A., Snyder-Mackler, N., (2021). Seasonal shifts in the gut microbiome indicate plastic responses to diet in wild geladas. Microbiome 9, 26. 10.1186/s40168-020-00977-9

Bates, D., Maechler, M., Bolker, B., Walker, S., Christensen, R. H. B., Singmann, H., … & Green, P. (2009). Package ‘lme4’. *URL* http://lme4.r-forge.r-project.org.

Bär, J., Leung, J.M., Hansen, C., Loke, P., Hall, A.R., Conour, L., Graham, A.L., (2020). Strong effects of lab-to-field environmental transitions on the bacterial intestinal microbiota of Mus musculus are modulated by Trichuris murisinfection. FEMS Microbiology Ecology 96, fiaa167. 10.1093/femsec/fiaa167

Bestion, E., Jacob, S., Zinger, L., Di Gesu, L., Richard, M., White, J., Cote, J., (2017). Climate warming reduces gut microbiota diversity in a vertebrate ectotherm. Nature Ecology & Evolution 1, 1–3. 10.1038/s41559-017-0161

Bibby, C. J. (1999). Making the most of birds as environmental indicators. Ostrich, 70(1), 81–88.

Bodawatta, K.H., Freiberga, I., Puzejova, K., Sam, K., Poulsen, M., Jønsson, K.A., (2021a). Flexibility and resilience of great tit (Parus major) gut microbiomes to changing diets. BMC Animal microbiome 3, 20. 10.1186/s42523-021-00076-6

Bodawatta, K.H., Hird, S.M., Grond, K., Poulsen, M., Jønsson, K.A., (2022a). Avian gut microbiomes taking flight. Trends in Microbiology 30, 268–280. 10.1016/j.tim.2021.07.003

Bodawatta, K.H., Klečková, I., Klečka, J., Pužejová, K., Koane, B., Poulsen, M., Jønsson, K.A., Sam, K., (2022b). Specific gut bacterial responses to natural diets of tropical birds. Science Reports 12, 713. 10.1038/s41598-022-04808-9

Bodawatta, K.H., Koane, B., Maiah, G., Sam, K., Poulsen, M., Jønsson, K.A., (2021b). Species-specific but not phylosymbiotic gut microbiomes of New Guinean passerine birds are shaped by diet and flight-associated gut modifications. Proceedings of the Royal. Society B 288, rspb.2021.0446, 20210446. 10.1098/rspb.2021.0446

Bolyen, E., Rideout, J.R., Dillon, M.R., Bokulich, N.A., Abnet, C., Al-Ghalith, G.A., Alexander, H., Alm, E.J., Arumugam, M., Asnicar, F., Bai, Y., Bisanz, J.E., Bittinger, K., Brejnrod, A., Brislawn, C.J., Brown, C.T., Callahan, B.J., Caraballo-Rodríguez, A.M., … Caporaso, J.G., (2018). QIIME 2: Reproducible, interactive, scalable, and extensible microbiome data science (preprint). PeerJ Preprints. 10.7287/peerj.preprints.27295v2

Bray, J.R., Curtis, J.T., (1957). An Ordination of the Upland Forest Communities of Southern Wisconsin. Ecological Monographs 27, 326–349. 10.2307/1942268

Brittingham, M.C., Temple, S.A., (1988). Impacts of Supplemental Feeding on Survival Rates of Black-Capped Chickadees. Ecology 69, 581–589. 10.2307/1941007

Callahan, B.J., McMurdie, P.J., Rosen, M.J., Han, A.W., Johnson, A.J.A., Holmes, S.P., (2016). DADA2: High-resolution sample inference from Illumina amplicon data. Nature Methods 13, 581–583. 10.1038/nmeth.3869

Cameron, E. S., Schmidt, P. J., Tremblay, B. J. M., Emelko, M. B., & Müller, K. M. (2020). To rarefy or not to rarefy: Enhancing diversity analysis of microbial communities through next-generation sequencing and rarefying repeatedly. BioRxiv, 2020-09.

Capunitan, D.C., Johnson, O., Terrill, R.S., Hird, S.M., (2020). Evolutionary signal in the gut microbiomes of 74 bird species from Equatorial Guinea. Molecular Ecology 29, 829–847. 10.1111/mec.15354

Chao, A., (2006). Species Estimation and Applications, in: Kotz, S., Read, C.B., Balakrishnan, N., Vidakovic, B., Johnson, N.L. (Eds.), Encyclopedia of Statistical Sciences. John Wiley & Sons, Inc., p. ess5051. 10.1002/0471667196.ess5051

Charmantier, A., McCleery, R.H., Cole, L.R., Perrins, C., Kruuk, L.E.B., Sheldon, B.C., (2008). Adaptive Phenotypic Plasticity in Response to Climate Change in a Wild Bird Population. Science 320, 800–803. 10.1126/science.1157174

Chevalier, C., Stojanović, O., Colin, D.J., Suarez-Zamorano, N., Tarallo, V., Veyrat-Durebex, C., Rigo, D., Fabbiano, S., Stevanović, A., Hagemann, S., Montet, X., Seimbille, Y., Zamboni, N., Hapfelmeier, S., Trajkovski, M., (2015). Gut Microbiota Orchestrates Energy Homeostasis during Cold. Cell 163, 1360– 1374. 10.1016/j.cell.2015.11.004

Cheville, N.F., Arp, L.H., (1978). Comparative pathologic findings of Escherichia coli infection in birds. Journal of the American Veterinary Medicine Association 173, 584–587.

Cody, M.L., (1981). Habitat Selection in Birds: The Roles of Vegetation Structure, Competitors, and Productivity. BioScience 31, 107–113. 10.2307/1308252

Corl, A., Charter, M., Rozman, G., Toledo, S., Turjeman, S., Kamath, P.L., Getz, W.M., Nathan, R., Bowie, R.C.K., (2020). Movement ecology and sex are linked to barn owl microbial community composition. Molecular Ecology 29, 1358–1371. 10.1111/mec.15398

Cox, A.R., Robertson, R.J., Lendvai, Á.Z., Everitt, K., Bonier, F., (2019). Rainy springs linked to poor nestling growth in a declining avian aerial insectivore (Tachycineta bicolor). Proceedings of the Royal Society B: Biological Sciences 286, 20190018. 10.1098/rspb.2019.0018

Davenport, E.R., Mizrahi-Man, O., Michelini, K., Barreiro, L.B., Ober, C., Gilad, Y., (2014). Seasonal Variation in Human Gut Microbiome Composition. Plos One 9, e90731. 10.1371/journal.pone.0090731

Davidson, G.L., Somers, S.E., Wiley, N., Johnson, C.N., Reichert, M.S., Ross, R.P., Stanton, C., Quinn, J.L., (2021). A time-lagged association between the gut microbiome, nestling weight and nestling survival in wild great tits. Journal of Animal Ecology 90, 989–1003. 10.1111/1365-2656.13428

Davis, N.M., Proctor, D.M., Holmes, S.P., Relman, D.A., Callahan, B.J., (2018). Simple statistical identification and removal of contaminant sequences in marker-gene and metagenomics data. Microbiome 6, 226. 10.1186/s40168-018-0605-2

Dietz, M. W., Matson, K. D., Versteegh, M. A., van der Velde, M., Parmentier, H. K., Arts, J. A., Salles, J. F. & Tieleman, B. I. (2022). Gut microbiota of homing pigeons shows summer–winter variation under constant diet indicating a substantial effect of temperature. Animal microbiome, 4(1), 64.

Dingemanse, N.J., Bouwman, K.M., van de Pol, M., van Overveld, T., Patrick, S.C., Matthysen, E., Quinn, J.L., (2012). Variation in personality and behavioural plasticity across four populations of the great tit Parus major. Journal of Animal Ecology 81, 116–126. 10.1111/j.1365-2656.2011.01877.x

Drobniak, S.M., Cichoń, M., Janas, K., Barczyk, J., Gustafsson, L., Zagalska-Neubauer, M., (2022). Habitat shapes diversity of gut microbiomes in a wild population of blue tits Cyanistes caeruleus. Journal of Avian Biology 2022, e02829. 10.1111/jav.02829

Du Toit, A., (2019). The gut microbiome and mental health. Nature Reviews Microbiology 17, 196–196. 10.1038/s41579-019-0163-z

Eren, A.M., Maignien, L., Sul, W.J., Murphy, L.G., Grim, S.L., Morrison, H.G., Sogin, M.L., Freckleton, R., (2013). Oligotyping: differentiating between closely related microbial taxa using 16S rRNA gene data. Methods in Ecology and Evolution 4, 1111–1119. 10.1111/2041-210X.12114

Escallón, C., Belden, L.K., Moore, I.T., (2019). The Cloacal Microbiome Changes with the Breeding Season in a Wild Bird. Integrative Organismal Biology 1, oby009. 10.1093/iob/oby009

Fontaine, S.S., Novarro, A.J., Kohl, K.D., (2018). Environmental temperature alters the digestive performance and gut microbiota of a terrestrial amphibian. Journal of Experimental Biology 221, jeb187559. 10.1242/jeb.187559

Fox, J., Weisberg, S., Adler, D., Bates, D., Baud-Bovy, G., Ellison, S., … & Heiberger, R. (2012). Package ‘car’. Vienna: R Foundation for Statistical Computing, 16.

Gadau, A., Crawford, M.S., Mayek, R., Giraudeau, M., McGraw, K.J., Whisner, C.M., Kondrat-Smith, C., Sweazea, K.L., (2019). A comparison of the nutritional physiology and gut microbiome of urban and rural house sparrows (Passer domesticus). Comparative Biochemistry and Physiology Part B: Biochemistry and Molecular Biology 237, 110332. 10.1016/j.cbpb.2019.110332

Gamero, A., Senar, J.C., Hohtola, E., Nilsson, J.-Å., Broggi, J., (2015). Population differences in the structure and coloration of great tit contour feathers: Feather colour and Microstructure. Biological Journal of the Linnean Society London 114, 82–91. 10.1111/bij.12409

Gilbert, J.A., Blaser, M.J., Caporaso, J.G., Jansson, J.K., Lynch, S.V., Knight, R., (2018). Current understanding of the human microbiome. Nature Medicine 24, 392–400. 10.1038/nm.4517

Góngora, E., Elliott, K.H., Whyte, L., (2021). Gut microbiome is affected by inter-sexual and inter-seasonal variation in diet for thick-billed murres (Uria lomvia). Science Reports 11, 1200. 10.1038/s41598-020-80557-x

Gregory, R.D., van Strien, A., Vorisek, P., Gmelig Meyling, A.W., Noble, D.G., Foppen, R.P.B., Gibbons, D.W., (2005). Developing indicators for European birds. Philosophical Transactions of the Royal Society B: Biological Sciences 360, 269–288. 10.1098/rstb.2004.1602

Grond, K., Sandercock, B.K., Jumpponen, A., Zeglin, L.H., (2018). The avian gut microbiota: community, physiology and function in wild birds. Journal of Avian Biology 49, e01788. 10.1111/jav.01788

Grubb, T.C., Jr., (1978). Weather-dependent Foraging Rates of Wintering Woodland Birds. The Auk 95, 370–376. 10.1093/auk/95.2.370

Haegeman, B., Hamelin, J., Moriarty, J., Neal, P., Dushoff, J., Weitz, J.S., (2013). Robust estimation of microbial diversity in theory and in practice. ISME Journal 7, 1092–1101. 10.1038/ismej.2013.10

Hartig, F., & Hartig, M. F. (2017). Package ‘DHARMa’. R package.

Hird, S.M., (2017). Evolutionary Biology Needs Wild Microbiomes. Fronters in Microbiology. 8, 725. 10.3389/fmicb.2017.00725

Hird, S.M., Sánchez, C., Carstens, B.C., Brumfield, R.T., (2015). Comparative Gut Microbiota of 59 Neotropical Bird Species. Frontiers in Microbiology 6.

Husby, A., Nussey, D.H., Visser, M.E., Wilson, A.J., Sheldon, B.C., Kruuk, L.E.B., (2010). Contrasting patterns of phenotypic plasticity in reproductive traits in two great tit (parus major) populations: multivariate patterns of phenotypic plasticity. Evolution. 10.1111/j.1558-5646.2010.00991.x

Ingala, M.R., Albert, L., Addesso, A., Watkins, M.J., Knutie, S.A., (2021). Differential effects of elevated nest temperature and parasitism on the gut microbiota of wild avian hosts. Animal microbiome 3, 67. 10.1186/s42523-021-00130-3

Jašarević, E., Morrison, K.E., Bale, T.L., (2016). Sex differences in the gut microbiome–brain axis across the lifespan. Philosophical Transactions of the Royal Society B: Biological Sciences 371, 20150122. 10.1098/rstb.2015.0122

Karr, J.R., (1976). Seasonality, Resource Availability, and Community Diversity in Tropical Bird Communities. The American Naturalist 110, 973–994. 10.1086/283121

Knutie, S.A., (2020). Food supplementation affects gut microbiota and immunological resistance to parasites in a wild bird species. Journal of Applied Ecology 57, 536–547. 10.1111/1365-2664.13567

Knutie, S.A., Chaves, J.A., Gotanda, K.M., (2019). Human activity can influence the gut microbiota of Darwin’s finches in the Galapagos Islands. Molecular Ecology 28, 2441–2450. 10.1111/mec.15088

Knutie, S.A., Gotanda, K.M., (2018). A Non-invasive Method to Collect Fecal Samples from Wild Birds for Microbiome Studies. Microbial Ecology 76, 851– 855. 10.1007/s00248-018-1182-4

Kohl, K.D., (2012). Diversity and function of the avian gut microbiota. Journal of Comparative Physiology B 182, 591–602. 10.1007/s00360-012-0645-z

Kohl, K.D., Yahn, J., (2016). Effects of environmental temperature on the gut microbial communities of tadpoles. Environmental Microbiology 18, 1561–1565. 10.1111/1462-2920.13255

Kopac, S.M., Klassen, J.L., (2016). Can They Make It on Their Own? Hosts, Microbes, and the Holobiont Niche. Frontiers in Microbiology 7.

Koskimies, P., (1989). Birds as a tool in environmental monitoring. Annales Zoologici Fennici 26, 153–166.

Krebs, J.R., (1971). Territory and Breeding Density in the Great Tit, Parus Major L. Ecology 52, 2–22. 10.2307/1934734

Kreisinger, J., Čížková, D., Kropáčková, L., Albrecht, T., (2015). Cloacal Microbiome Structure in a Long-Distance Migratory Bird Assessed Using Deep 16sRNA Pyrosequencing. Plos one 10, e0137401. 10.1371/journal.pone.0137401

Lahti, L., & Shetty, S. (2018). Introduction to the microbiome R package. *Preprint at* https://microbiome.github.io/tutorials.

Lemoine, M., Lucek, K., Perrier, C., Saladin, V., Adriaensen, F., Barba, E., Belda, E.J., Charmantier, A., Cichoń, M., Eeva, T., Grégoire, A., Hinde, C.A., Johnsen, A., Komdeur, J., Mänd, R., Matthysen, E., Norte, A.C., Pitala, N., Sheldon, B.C., … Richner, H., (2016). Low but contrasting neutral genetic differentiation shaped by winter temperature in European great tits. Biological Journal of the Linnean Society 118, 668–685. 10.1111/bij.12745

Lewis, W.B., Moore, F.R., Wang, S., (2017). Changes in gut microbiota of migratory passerines during stopover after crossing an ecological barrier. The Auk 134, 137–145. 10.1642/AUK-16-120.1

Ley, R.E., Hamady, M., Lozupone, C., Turnbaugh, P.J., Ramey, R.R., Bircher, J.S., Schlegel, M.L., Tucker, T.A., Schrenzel, M.D., Knight, R., Gordon, J.I., (2008). Evolution of Mammals and Their Gut Microbes. Science 320, 1647–1651. 10.1126/science.1155725

Liu, H., Guo, X., Gooneratne, R., Lai, R., Zeng, C., Zhan, F., Wang, W., (2016). The gut microbiome and degradation enzyme activity of wild freshwater fishes influenced by their trophic levels. Science Reports 6, 24340. 10.1038/srep24340

Liukkonen, M., Hukkanen, M., Cossin-Sevrin, N., Stier, A., Vesterinen, E., Grond, K., Ruuskanen, S., (2023). No evidence for associations between brood size, gut microbiome diversity and survival in great tit (Parus major) nestlings. Animal microbiome 5, 19. 10.1186/s42523-023-00241-z

Loo, W.T., García-Loor, J., Dudaniec, R.Y., Kleindorfer, S., Cavanaugh, C.M., (2019). Host phylogeny, diet, and habitat differentiate the gut microbiomes of Darwin’s finches on Santa Cruz Island. Science Reports 9, 18781. 10.1038/s41598-019-54869-6

Love, M., Anders, S., & Huber, W. (2014). Differential analysis of count data–the DESeq2 package. Genome Biology, 15(550), 10–1186.

Lucas, G., (2018). Gut thinking: the gut microbiome and mental health beyond the head. Microbial Ecology in Health and Disease 29, 1548250. 10.1080/16512235.2018.1548250

Martin, M., (2011). Cutadapt removes adapter sequences from high-throughput sequencing reads. EMBnet.journal 17, 10–12. 10.14806/ej.17.1.200

Maurice, C.F., CL Knowles, S., Ladau, J., Pollard, K.S., Fenton, A., Pedersen, A.B., Turnbaugh, P.J., (2015). Marked seasonal variation in the wild mouse gut microbiota. ISME Journal 9, 2423–2434. 10.1038/ismej.2015.53

McKenzie, C., Tan, J., Macia, L., Mackay, C.R., (2017). The nutrition-gut microbiome-physiology axis and allergic diseases. Immunological Reviews 278, 277–295. 10.1111/imr.12556

McMurdie, P.J., Holmes, S., (2013). phyloseq: An R Package for Reproducible Interactive Analysis and Graphics of Microbiome Census Data. Plos One 8, e61217. 10.1371/journal.pone.0061217

McNamara, J.M., Houston, A.I., Lima, S.L., (1994). Foraging Routines of Small Birds in Winter: A Theoretical Investigation. Journal of Avian Biology 25, 287–302. 10.2307/3677276

Meng, H., Zhang, Y., Zhao, L., Zhao, W., He, C., Honaker, C.F., Zhai, Z., Sun, Z., Siegel, P.B., (2014). Body Weight Selection Affects Quantitative Genetic Correlated Responses in Gut Microbiota. Plos One 9, e89862. 10.1371/journal.pone.0089862

Moeller, A.H., Ivey, K., Cornwall, M.B., Herr, K., Rede, J., Taylor, E.N., Gunderson, A.R., (2020). The Lizard Gut Microbiome Changes with Temperature and Is Associated with Heat Tolerance. Applied and Environmental Microbiology 86, e01181–20. 10.1128/AEM.01181-20

Murray, M.H., Lankau, E.W., Kidd, A.D., Welch, C.N., Ellison, T., Adams, H.C., Lipp, E.K., Hernandez, S.M., (2020). Gut microbiome shifts with urbanization and potentially facilitates a zoonotic pathogen in a wading bird. Plos One 15, e0220926. 10.1371/journal.pone.0220926

Naef-Daenzer, B., Grüebler, M.U., (2008). Post-Fledging Range use of Great Tit Parus major Families in Relation to Chick Body Condition. Ardea 96, 181–190. 10.5253/078.096.0204

Naef-Daenzer, B., Widmer, F., Nuber, M., (2001). Differential post-fledging survival of great and coal tits in relation to their condition and fledging date. Journal of Animal Ecology 70, 730–738. 10.1046/j.0021-8790.2001.00533.x

Noguera, J.C., Aira, M., Pérez-Losada, M., Domínguez, J., Velando, A., (2018). Glucocorticoids modulate gastrointestinal microbiome in a wild bird. Royal Society Open Science 5, 171743. 10.1098/rsos.171743

Noordwijk, A.J.V., Van Balen, J.H., Scharloo, W., (2002). Heritability of Ecologically Important Traits in the Great Tit. Ardea 38–90, 193–203. 10.5253/arde.v68.p193

Norte, A.C., Ramos, J.A., Sampaio, H.L., Sousa, J.P., Sheldon, B.C., (2010). Physiological Condition and Breeding Performance of the Great Tit. The Condor 112, 79–86. 10.1525/cond.2010.08007

Oksanen, J., Blanchet, F. G., Friendly, M., Kindt, R., Legendre, P., McGlinn, D., … & Imports, M. A. S. S. (2019). Package ‘vegan’. Community ecology package, version, 2(9).

Parada, A.E., Needham, D.M., Fuhrman, J.A., (2016). Every base matters: assessing small subunit rRNA primers for marine microbiomes with mock communities, time series and global field samples. Environmental Microbiology 18, 1403–1414. 10.1111/1462-2920.13023

Pereira, H.M., Cooper, H.D., (2006). Towards the global monitoring of biodiversity change. Trends in Ecology & Evolution 21, 123–129. 10.1016/j.tree.2005.10.015

Phillips, J.N., Berlow, M., Derryberry, E.P., (2018). The Effects of Landscape Urbanization on the Gut Microbiome: An Exploration Into the Gut of Urban and Rural White-Crowned Sparrows. Frontiers in Ecology and Evolution 6, 148. 10.3389/fevo.2018.00148

Pigot, A.L., Sheard, C., Miller, E.T., Bregman, T.P., Freeman, B.G., Roll, U., Seddon, N., Trisos, C.H., Weeks, B.C., Tobias, J.A., (2020). Macroevolutionary convergence connects morphological form to ecological function in birds. Nature Ecology & Evolution 4, 230–239. 10.1038/s41559-019-1070-4

Quast, C., Pruesse, E., Yilmaz, P., Gerken, J., Schweer, T., Yarza, P., Peplies, J., Glöckner, F.O., (2013). The SILVA ribosomal RNA gene database project: improved data processing and web-based tools. Nucleic Acids Research 41, D590–D596. 10.1093/nar/gks1219

Radford, A.N., McCleery, R.H., Woodburn, R.J.W., Morecroft, M.D., (2001). Activity patterns of parent Great Tits Parus major feeding their young during rainfall. Bird Study 48, 214–220. 10.1080/00063650109461220

Rahbek, C., Graves, G.R., (2001). Multiscale assessment of patterns of avian species richness. Proceedings of the National Academy of Sciences 98, 4534–4539. 10.1073/pnas.071034898

Reid, G., Burton, J., (2002). Use of Lactobacillus to prevent infection by pathogenic bacteria. Microbes and Infection 4, 319–324. 10.1016/S1286-4579(02)01544-7

Ren, T., Boutin, S., Humphries, M.M., Dantzer, B., Gorrell, J.C., Coltman, D.W., McAdam, A.G., Wu, M., (2017). Seasonal, spatial, and maternal effects on gut microbiome in wild red squirrels. Microbiome 5, 163. 10.1186/s40168-017-0382-3

Rosenberg, E., Zilber-Rosenberg, I., (2018). The hologenome concept of evolution after 10 years. Microbiome 6, 78. 10.1186/s40168-018-0457-9

Rosenberg, E., & Zilber-Rosenberg, I. (2016). Do microbiotas warm their hosts?. Gut microbes, 7(4), 283–285.

Rothschild, D., Weissbrod, O., Barkan, E., Kurilshikov, A., Korem, T., Zeevi, D., Costea, P.I., Godneva, A., Kalka, I.N., Bar, N., Shilo, S., Lador, D., Vila, A.V., Zmora, N., Pevsner-Fischer, M., Israeli, D., Kosower, N., Malka, G., Wolf, B.C., … & Segal, E., 2018. Environment dominates over host genetics in shaping human gut microbiota. Nature 555, 210–215. 10.1038/nature25973

Sabree, Z.L., Moran, N.A., (2014). Host-specific assemblages typify gut microbial communities of related insect species. Springer Plus 3, 138. 10.1186/2193-1801-3-138

Saulnier, A., Bleu, J., Boos, A., Millet, M., Zahn, S., Ronot, P., El Masoudi, I., Rojas, E.R., Uhlrich, P., Del Nero, M., Massemin, S., (2023). Inter-annual variation of physiological traits between urban and forest great tits. Comparative Biochemistry and Physiology Part A: Molecular & Integrative Physiology 279, 111385. 10.1016/j.cbpa.2023.111385

Schloss, P.D., (2023). Waste not, want not: Revisiting the analysis that called into question the practice of rarefaction. BioRxiv 2023-06. 10.1101/2023.06.23.546312

Schmiedová, L., Kreisinger, J., Kubovčiak, J., Těšický, M., Martin, J.-F., Tomášek, O., Kauzálová, T., Sedláček, O., Albrecht, T., (2023). Gut microbiota variation between climatic zones and due to migration strategy in passerine birds. Frontiers in Microbiology 14.

Schmiedová, L., Tomášek, O., Pinkasová, H., Albrecht, T., Kreisinger, J., (2022). Variation in diet composition and its relation to gut microbiota in a passerine bird. Science Reports 12, 3787. 10.1038/s41598-022-07672-9

Schöll, E.M., Ohm, J., Hoffmann, K.F., Hille, S.M., (2016). Caterpillar biomass depends on temperature and precipitation, but does not affect bird reproduction. Acta Oecologica 74, 28–36. 10.1016/j.actao.2016.06.004

Sepulveda, J., Moeller, A.H., (2020). The Effects of Temperature on Animal Gut Microbiomes. Frontiers in Microbiology 11.

Singh, R.K., Chang, H.-W., Yan, D., Lee, K.M., Ucmak, D., Wong, K., Abrouk, M., Farahnik, B., Nakamura, M., Zhu, T.H., Bhutani, T., Liao, W., (2017). Influence of diet on the gut microbiome and implications for human health. Journal of Translational Medicine 15, 73. 10.1186/s12967-017-1175-y

Spor, A., Koren, O., Ley, R., (2011). Unravelling the effects of the environment and host genotype on the gut microbiome. Nature Reviews Microbiology 9, 279–290. 10.1038/nrmicro2540

Sullam, K.E., Essinger, S.D., Lozupone, C.A., O’connor, M.P., Rosen, G.L., Knight, R., Kilham, S.S., Russell, J.A., (2012). Environmental and ecological factors that shape the gut bacterial communities of fish: a meta-analysis. Molecular Ecology 21, 3363–3378. 10.1111/j.1365-294X.2012.05552.x

Sullam, K.E., Rubin, B.E., Dalton, C.M., Kilham, S.S., Flecker, A.S., Russell, J.A., (2015). Divergence across diet, time and populations rules out parallel evolution in the gut microbiomes of Trinidadian guppies. ISME Journal 9, 1508–1522. 10.1038/ismej.2014.231

Tajima, K., Nonaka, I., Higuchi, K., Takusari, N., Kurihara, M., Takenaka, A., Mitsumori, M., Kajikawa, H., Aminov, R.I., (2007). Influence of high temperature and humidity on rumen bacterial diversity in Holstein heifers. Anaerobe 13, 57–64. 10.1016/j.anaerobe.2006.12.001

Teyssier, A., Lens, L., Matthysen, E., White, J., (2018a). Dynamics of Gut Microbiota Diversity During the Early Development of an Avian Host: Evidence From a Cross-Foster Experiment. Frontiers in Microbiology 9, 1524. 10.3389/fmicb.2018.01524

Teyssier, A., Matthysen, E., Hudin, N.S., De Neve, L., White, J., Lens, L., (2020). Diet contributes to urban-induced alterations in gut microbiota: experimental evidence from a wild passerine. Proceedings of the Royal Society: Biology 287, 20192182. 10.1098/rspb.2019.2182

Teyssier, A., Rouffaer, L.O., Saleh Hudin, N., Strubbe, D., Matthysen, E., Lens, L., White, J., (2018b). Inside the guts of the city: Urban-induced alterations of the gut microbiota in a wild passerine. Science of The Total Environment 612, 1276– 1286. 10.1016/j.scitotenv.2017.09.035

Thie, N., Corl, A., Turjeman, S., Efrat, R., Kamath, P.L., Getz, W.M., Bowie, R.C.K., Nathan, R., (2022). Linking migration and microbiota at a major stopover site in a long-distance avian migrant. Movement Ecology 10, 46. 10.1186/s40462-022-00347-0

Tian, Y., Li, G., Chen, L., Bu, X., Shen, J., Tao, Z., Zeng, T., Du, X., Lu, L., (2020). High-temperature exposure alters the community structure and functional features of the intestinal microbiota in Shaoxing ducks (Anas platyrhynchos). Poultry Science 99, 2662–2674. 10.1016/j.psj.2019.12.046

Tilg, H., Kaser, A., (2011). Gut microbiome, obesity, and metabolic dysfunction. Journal of Clinical Investigation 121, 2126–2132. 10.1172/JCI58109

Tinya, F., Kovács, B., Bidló, A., Dima, B., Király, I., Kutszegi, G., Lakatos, F., Mag, Z., Márialigeti, S., Nascimbene, J., Samu, F., Siller, I., Szél, G., Ódor, P., (2021). Environmental drivers of forest biodiversity in temperate mixed forests – A multi-taxon approach. Science of The Total Environment 795, 148720. 10.1016/j.scitotenv.2021.148720

Tizard, I., (2004). Salmonellosis in wild birds. Seminars in Avian and Exotic Pet Medicine, Emerging Diseases 13, 50–66. 10.1053/j.saep.2004.01.008

Tong, Q., Cui, L.-Y., Hu, Z.-F., Du, X.-P., Abid, H.M., Wang, H.-B., (2020). Environmental and host factors shaping the gut microbiota diversity of brown frog Rana dybowskii. Science of The Total Environment 741, 140142. 10.1016/j.scitotenv.2020.140142

Vel’ký, M., Kaňuch, P., Krištín, A., (2011). Food composition of wintering great tits (Parus major): habitat and seasonal aspects. Folia Zoologica 60, 228–236. 10.25225/fozo.v60.i3.a7.2011

Waite, D.W., Taylor, M.W., (2014). Characterizing the avian gut microbiota: membership, driving influences, and potential function. Frontiers in Microbiology 5.

Wang, Feng, J.H., Zhang, M.H., Li, X.M., Ma, D.D., Chang, S.S., (2018). Effects of high ambient temperature on the community structure and composition of ileal microbiome of broilers. Poultry Science 97, 2153–2158. 10.3382/ps/pey032

Wang, S., Chen, L., He, M., Shen, J., Li, G., Tao, Z., Wu, R., Lu, L., (2018). Different rearing conditions alter gut microbiota composition and host physiology in Shaoxing ducks. Science Reports 8, 7387. 10.1038/s41598-018-25760-7

Weinroth, M.D., Belk, A.D., Dean, C., Noyes, N., Dittoe, D.K., Rothrock, M.J., Jr, Ricke, S.C., Myer, P.R., Henniger, M.T., Ramírez, G.A., Oakley, B.B., Summers, K.L., Miles, A.M., Ault-Seay, T.B., Yu, Z., Metcalf, J.L., Wells, J.E., (2022). Considerations and best practices in animal science 16S ribosomal RNA gene sequencing microbiome studies. Journal of Animal Science 100, skab346. 10.1093/jas/skab346

Wen, C., Yan, W., Mai, C., Duan, Z., Zheng, J., Sun, C., Yang, N., (2021). Joint contributions of the gut microbiota and host genetics to feed efficiency in chickens. Microbiome 9, 126. 10.1186/s40168-021-01040-x

Williams, C.M., Henry, H.A.L., Sinclair, B.J., (2015). Cold truths: how winter drives responses of terrestrial organisms to climate change. Biological Reviews 90, 214–235. 10.1111/brv.12105

Worsley, S.F., Davies, C.S., Mannarelli, M.-E., Hutchings, M.I., Komdeur, J., Burke, T., Dugdale, H.L., Richardson, D.S., (2021). Gut microbiome composition, not alpha diversity, is associated with survival in a natural vertebrate population. Animal microbiome 3, 84. 10.1186/s42523-021-00149-6

Worthmann, A., John, C., Rühlemann, M.C., Baguhl, M., Heinsen, F.-A., Schaltenberg, N., Heine, M., Schlein, C., Evangelakos, I., Mineo, C., Fischer, M., Dandri, M., Kremoser, C., Scheja, L., Franke, A., Shaul, P.W., Heeren, J., (2017). Cold-induced conversion of cholesterol to bile acids in mice shapes the gut microbiome and promotes adaptive thermogenesis. Nature Medicine 23, 839–849. 10.1038/nm.4357

Xiao, G., Liu, S., Xiao, Y., Zhu, Y., Zhao, H., Li, A., Li, Z., Feng, J., (2019). Seasonal Changes in Gut Microbiota Diversity and Composition in the Greater Horseshoe Bat. Frontiers in Microbiology 10.

Yang, Y., Gao, H., Li, X., Cao, Z., Li, M., Liu, J., Qiao, Y., Ma, L., Zhao, Z., Pan, H., (2021). Correlation analysis of muscle amino acid deposition and gut microbiota profile of broilers reared at different ambient temperatures. Animal Bioscience 34, 93–101. 10.5713/ajas.20.0314

Yilmaz, P., Parfrey, L.W., Yarza, P., Gerken, J., Pruesse, E., Quast, C., Schweer, T., Peplies, J., Ludwig, W., Glöckner, F.O., (2014). The SILVA and “All-species Living Tree Project (LTP)” taxonomic frameworks. Nucleic Acids Research 42, D643–D648. 10.1093/nar/gkt1209

Zhang, X.-Y., Sukhchuluun, G., Bo, T.-B., Chi, Q.-S., Yang, J.-J., Chen, B., Zhang, L., Wang, D.-H., (2018). Huddling remodels gut microbiota to reduce energy requirements in a small mammal species during cold exposure. Microbiome 6, 103. 10.1186/s40168-018-0473-9

Zhao, L., Wang, G., Siegel, P., He, C., Wang, H., Zhao, W., Zhai, Z., Tian, F., Zhao, J., Zhang, H., Sun, Z., Chen, W., Zhang, Y., Meng, H., (2013). Quantitative Genetic Background of the Host Influences Gut Microbiomes in Chickens. Science Reports 3, 1163. 10.1038/srep01163

Zheng, D., Liwinski, T., Elinav, E., (2020). Interaction between microbiota and immunity in health and disease. Cell Research 30, 492–506. 10.1038/s41422-020-0332-7

Zhu, L., Liao, R., Wu, N., Zhu, G., Yang, C., (2019). Heat stress mediates changes in fecal microbiome and functional pathways of laying hens. Applied Microbiology and Biotechnology 103, 461–472. 10.1007/s00253-018-9465-8

Zilber-Rosenberg, I., Rosenberg, E., (2008). Role of microorganisms in the evolution of animals and plants: the hologenome theory of evolution. FEMS Microbiology Reviews 32, 723–735. 10.1111/j.1574-6976.2008.00123.x

