## Supplementary file for "Seasonal and environmental factors contribute to the variation in the gut microbiome: a large-scale study of a small bird"

### SI 1: Sample numbers by population and season with details about regional average rainfall and temperature per month

| **Population** | **Winter** | **Summer** | **Winter rainfall range 2 weeks prior to sampling** | **Summer rainfall range**  **2 weeks prior to sampling** | **Winter temperature range**  **2 weeks prior to sampling** | **Summer temperature range**  **2 weeks prior to sampling** |
| --- | --- | --- | --- | --- | --- | --- |
| **Pilis-Visegrád Mountains , Hungary**  47.7236 N, 19.0225 E | 22 | 27 | 0.71-1.79 mm, February | 0.76-2.46 mm, May | -0.13 - 3.3 C, February | 13.99 - 16.8 C, May |
| **Jyväskylä, Finland**  62.1938 N, 25.7475 E | 27 | 25 | 0.31-2.64 mm, January / February | 0.14-5.51 mm, May / June | -15.5 - -6.99 C, January / February | 8.6 - 22.3 C, May / June |
| **Lund, Sweden**  55.7241 N, 13.2772 E | 26 | 21 | 0.97-1.76mm, February | 1.59 mm, June | -4.5 - 4.8 C, February | 8.7 - 14.5 C, June |
| **Oulu, Finland**  65.01687 N, 25.4236 E | 21 | 20 | 0.19-2.66 mm, January | 0.07-10.08 mm, June | -12.78 - -4.89 C, January | 13.98 – 16.46 C, June |
| **La Hiruela, Spain**  40.7615 N, 3.7925 W | 0 | 12 | - | 0.14-0.91 mm, June | - | 17.88 – 24.2 C, June |
| **Tartu, Estonia**  58.2077 N, 26.2721 E | 7 | 20 | 1.58-2.33 mm, February | 4.84-4.86 mm, May | -5.43 - -2.34 C, February | 10.4 – 11.6 C, May / June |
| **Turku, Finland**  60.4341 N, 22.1541 E | 21 | 16 | 1.79-2.75 mm, January | 0.33-2.34 mm, June | -6.07 - -5.68 C, January | 11.67 – 15.9 C, June |
| **Westerheide, Netherlands**  52.0056 N, 5.5013 E | 0 | 20 | - | 2.04-3.39 mm, May | - | 11.44 - 13.16 C, May |
| ***Total*** | *124* | *161* |  |  |  |  |

| **Location** | **Habitat** | **Winter diet** |
| --- | --- | --- |
| **Pilis-Visegrád Mountains, Hungary**  47.7236 N, 19.0225 E | Deciduous | Sunflower seeds |
| **Jyväskylä, Finland**  62.1938 N, 25.7475 E | Mixed coniferous and deciduous | Peanuts & sunflower seeds |
| **Lund, Sweden**  55.7241 N, 13.2772 E | Deciduous | Sunflower seeds |
| **Oulu, Finland**  65.01687 N, 25.4236 E | Deciduous | Sunflower seeds |
| **La Hiruela, Spain**  40.7615 N, 3.7925 W | Mixed coniferous and deciduous | - |
| **Tartu, Estonia**  58.2077 N, 26.2721 E | Mixed coniferous and deciduous | Sunflower seeds |
| **Turku, Finland**  60.4341 N, 22.1541 E | Mixed coniferous and deciduous | Peanuts and sunflower seeds |
| **Westerheide, Netherlands**  52.0056 N, 5.5013 E | Deciduous | - |

### SI 2: Sequence summaries for unrarefied and rarefied data and the number of contaminants removed

| **Dataset** | **Number of samples** | **Number of taxa** | **Min. reads** | **Max. reads** | **Average reads** | **Total reads** | **Median reads** |
| --- | --- | --- | --- | --- | --- | --- | --- |
| **Unrarefied** | 284:  124 winter and 161 summer | 15 288 | 107 | 532 664 | 57 740.70 | 16 629 323 | 17 189 |
| **Rarefied (1000 taxa)** | 277:  121 winter and 156 summer | 6883 | 1000 | 1000 | 1000 | 280 000 | 1000 |
| **Removed contaminants** | **-** | 61 | 339 | 368 693 | 36 668.32 | 806 703 | 7662 |

### SI 3: Gut microbiome relative abundance per sample

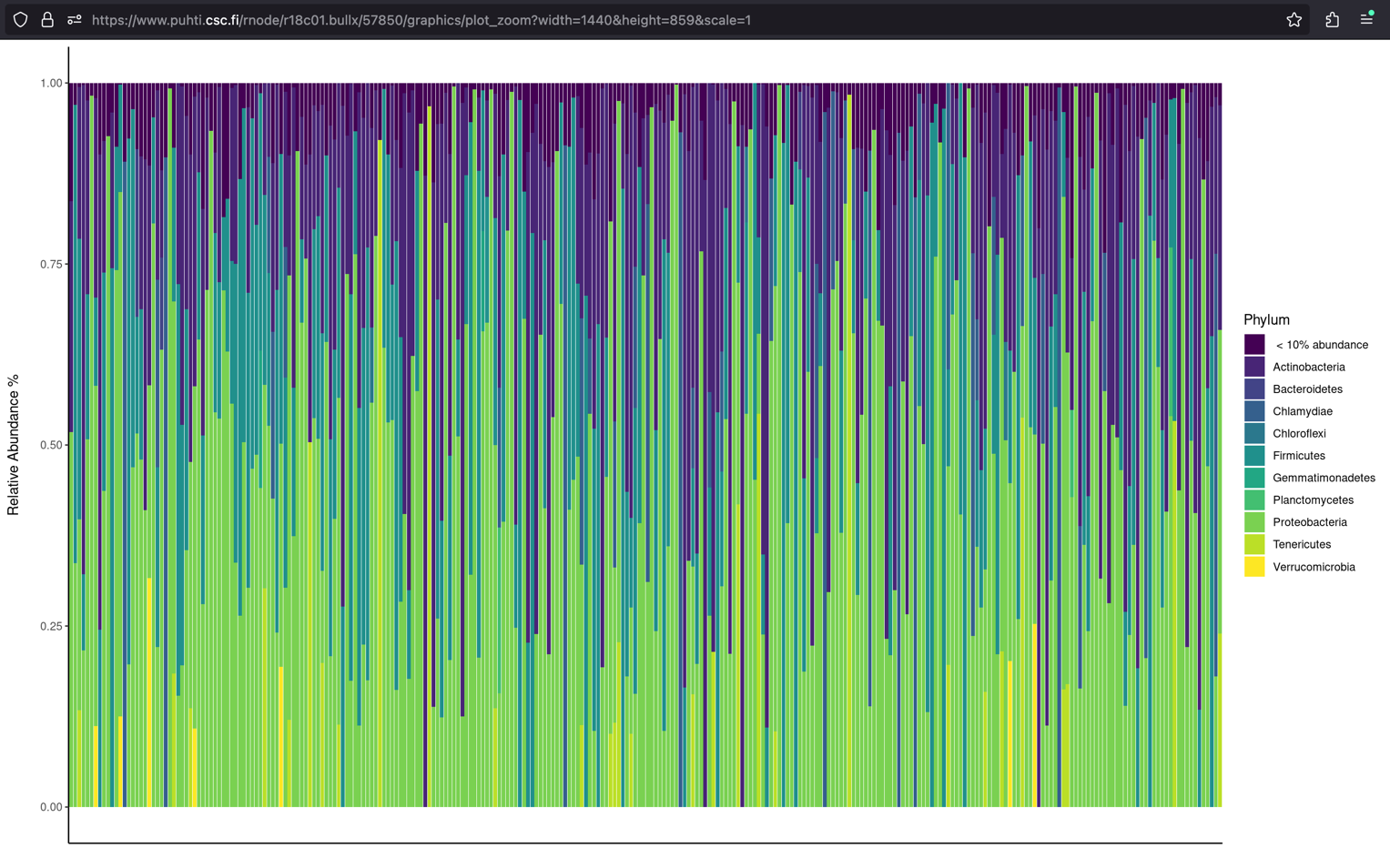

*Top 10 most abundant phyla are specified by color and taxa that are less than 10 % in total abundance are grouped together.

### SI 4. Linear model measuring which population specific factors contribute to gut microbiome alpha diversity.

In table 1) the linear model includes body condition as a physiological factor, and in table 2) body condition is replaced with weight to check consistency of results.

Section A) estimates for the linear model investigating how season and population associate with gut microbiome alpha diversity.

Section B) estimates for the linear model investigating how latitude, habitat, temperature, rainfall, sex, and body condition associate with gut microbiome alpha diversity (data across both seasons).

Section C) estimates for the linear model investigating how latitude, habitat, temperature, rainfall, diet, sex, and body condition associate with gut microbiome alpha diversity during winter.

Section D) estimates for the linear model investigating how latitude, habitat, temperature, rainfall, sex, and body condition associate with gut microbiome alpha diversity during summer.

**1)**

*Oulu as the northernmost population is used as the reference level in models in which population is used as an explanatory variable.

| **Shannon** | **estimate** | **s.e.** | **t** | **P** | **Chao1** | **estimate** | **s.e.** | **t** | **P** |
| --- | --- | --- | --- | --- | --- | --- | --- | --- | --- |
| **A** |  |  |  |  |  |  |  |  |  |
| **Season (winter)** | 0.385 | 0.149 | 2.575 | ***0.011*** |  | 32.423 | 25.493 | 1.272 | 0.205 |
| **Population Jyvaskyla** | -0.043 | 0.243 | -0.179 | 0.858 |  | 24.652 | 41.427 | 0.595 | 0.552 |
| **Population Turku** | 0.098 | 0.261 | 0.376 | 0.707 |  | -7.318 | 44.608 | -0.164 | 0.870 |
| **Population Tartu** | 0.403 | 0.294 | 1.367 | 0.173 |  | 51.834 | 50.244 | 1.032 | 0.303 |
| **Population Lund** | 0.052 | 0.248 | 0.210 | 0.834 |  | 35.922 | 42.274 | 0.850 | 0.396 |
| **Population Pilis-Visegrád Mountains** | 0.071 | 0.247 | 0.290 | 0.772 |  | -38.693 | 42.076 | 0.920 | 0.359 |
| **B** |  |  |  |  |  |  |  |  |  |
| ***Both seasons*** |  |  |  |  |  |  |  |  |  |
| **Latitude** | -0.026 | 0.015 | -1.799 | 0.073 |  | -2.068 | 2.479 | -0.834 | 0.405 |
| **Habitat (mixed)** | 0.195 | 0.164 | 1.193 | 0.234 |  | 11.279 | 27.745 | 0.407 | 0.685 |
| **Temperature** | -0.014 | 0.009 | -1.583 | 0.115 |  | -1.938 | 1.511 | -1.282 | 0.201 |
| **Rainfall** | -0.010 | 0.043 | -0.220 | 0.826 |  | -1.607 | 7.361 | -0.218 | 0.827 |
| **Sex (male)** | -0.067 | 0.157 | -0.423 | 0.673 |  | -36.515 | 26.684 | -1.368 | 0.173 |
| **Body condition** | 0.134 | 0.079 | 1.698 | 0.091 |  | 1.336 | 13.418 | 0.200 | 0.921 |
| **C** |  |  |  |  |  |  |  |  |  |
| ***Winter*** |  |  |  |  |  |  |  |  |  |
| **Latitude** | -0.014 | 0.036 | -0.383 | 0.703 |  | -7.649 | 6.703 | -1.141 | 0.257 |
| **Habitat (mixed)** | 1.020 | 0.449 | 2.274 | ***0.025*** |  | 64.634 | 83.329 | 0.776 | 0.440 |
| **Temperature** | 0.031 | 0.035 | 0.877 | 0.383 |  | 0.468 | 6.526 | 0.072 | 0.943 |
| **Rainfall** | -0.093 | 0.227 | -0.410 | 0.683 |  | 4.069 | 42.154 | 0.097 | 0.923 |
| **Diet (sunflower seeds)** | 0.485 | 0.444 | 1.092 | 0.277 |  | 83.383 | 82.482 | 1.011 | 0.315 |
| **Sex (male)** | 0.132 | 0.251 | 0.525 | 0.601 |  | -76.576 | 46.657 | -1.641 | 0.104 |
| **Body condition** | 0.186 | 0.115 | 1.617 | 0.109 |  | 23.050 | 21.353 | 1.079 | 0.283 |
| **D** |  |  |  |  |  |  |  |  |  |
| ***Summer*** |  |  |  |  |  |  |  |  |  |
| **Latitude** | -0.022 | 0.017 | -1.286 | 0.201 |  | 0.381 | 2.577 | 0.148 | 0.883 |
| **Habitat (mixed)** | 0.024 | 0.211 | 0.116 | 0.908 |  | 39.758 | 32.321 | 1.230 | 0.221 |
| **Temperature** | 0.009 | 0.037 | 0.251 | 0.802 |  | -5.741 | 5.654 | -1.015 | 0.312 |
| **Rainfall** | -0.007 | 0.046 | -0.160 | 0.873 |  | -3.490 | 7.080 | -0.493 | 0.623 |
| **Sex (male)** | -0.300 | 0.210 | -1.433 | 0.154 |  | -1.127 | 32.197 | -0.035 | 0.972 |
| **Body condition** | 0.150 | 0.125 | 1.201 | 0.232 |  | -6.739 | 19.518 | -0.345 | 0.730 |

**2)**

| **Shannon** | **estimate** | **s.e.** | **t** | **P** | **Chao1** | **estimate** | **s.e.** | **t** | **P** |
| --- | --- | --- | --- | --- | --- | --- | --- | --- | --- |
| **A** |  |  |  |  |  |  |  |  |  |
| ***Both seasons*** |  |  |  |  |  |  |  |  |  |
| **Latitude** | -0.025 | 0.014 | -1.809 | 0.072 |  | -0.760 | 2.365 | -0.321 | 0.748 |
| **Habitat (mixed)** | 0.160 | 0.149 | 1.071 | 0.258 |  | -4.421 | 25.293 | -0.175 | 0.861 |
| **Temperature** | -0.015 | 0.009 | -1.682 | 0.099 |  | -1.906 | 1.463 | -1.303 | 0.194 |
| **Rainfall** | -0.009 | 0.042 | -0.217 | 0.828 |  | -4.249 | 7.050 | -0.603 | 0.547 |
| **Sex (male)** | -0.178 | 0.150 | -1.182 | 0.238 |  | -29.547 | 25.502 | -1.159 | 0.248 |
| **Weight** | 0.118 | 0.073 | 1.609 | 0.109 |  | -2.000 | 12.377 | -0.162 | 0.872 |
| **B** |  |  |  |  |  |  |  |  |  |
| ***Winter*** |  |  |  |  |  |  |  |  |  |
| **Latitude** | -0.013 | 0.036 | -0.036 | 0.718 |  | -7.407 | 6.733 | -1.100 | 0.274 |
| **Habitat (mixed)** | 0.952 | 0.440 | 2.264 | ***0.033*** |  | 54.072 | 81.702 | 0.662 | 0.510 |
| **Temperature** | 0.027 | 0.035 | 0.781 | 0.437 |  | -0.126 | 6.468 | -0.019 | 0.985 |
| **Rainfall** | -0.106 | 0.228 | -0.464 | 0.644 |  | 2.594 | 42.245 | 0.061 | 0.951 |
| **Diet (sunflower seeds)** | 0.483 | 0.446 | 1.085 | 0.281 |  | 82.277 | 82.712 | 0.995 | 0.322 |
| **Sex (male)** | 0.048 | 0.266 | 0.179 | 0.859 |  | -85.277 | 49.404 | 0.995 | 0.322 |
| **Weight** | 0.159 | 0.108 | 1.468 | 0.145 |  | 17.782 | 20.055 | 0.997 | 0.378 |
| **C** |  |  |  |  |  |  |  |  |  |
| ***Summer*** |  |  |  |  |  |  |  |  |  |
| **Latitude** | -0.022 | 0.016 | -1.440 | 0.152 |  | 2.082 | 2.400 | 0.868 | 0.387 |
| **Habitat (mixed)** | 0.027 | 0.192 | 0.139 | 0.890 |  | 22.009 | 29.601 | 0.744 | 0.458 |
| **Temperature** | 0.004 | 0.033 | 0.128 | 0.898 |  | -1.666 | 5.008 | -0.333 | 0.740 |
| **Rainfall** | -0.005 | 0.044 | -0.106 | 0.916 |  | -4.872 | 6.807 | -0.716 | 0.475 |
| **Sex (male)** | -0.410 | 0.184 | -2.226 | 0.050 |  | 5.725 | 28.359 | 0.202 | 0.840 |
| **Weight** | 0.106 | 0.112 | 0.951 | 0.343 |  | -4.755 | 17.231 | -0.276 | 0.783 |

### SI 5. A linear model with interaction to test whether there is a significant interaction between season and population in relation to alpha diversity.

A) Estimates for the linear model
B) Type 3 analysis of variance for the linear model

**A)**

| **Shannon** | **estimate** | **s.e.** | **t** | **P** | **Chao1** | **Estimate** | **S.E.** | **t** | **P** |
| --- | --- | --- | --- | --- | --- | --- | --- | --- | --- |
| **Population** |  |  |  |  |  |  |  |  |  |
| **Jyvaskyla** | -0.205 | 0.350 | -0.587 | 0.558 |  | 65.092 | 59.212 | 1.099 | 0.273 |
| **Turku** | -0.175 | 0.388 | -0.451 | 0.652 |  | -5.325 | 65.596 | -0.081 | 0.935 |
| **Tartu** | 0.034 | 0.376 | 0.090 | 0.926 |  | 35.439 | 63.539 | 0.558 | 0.578 |
| **Lund** | -0.004 | 0.370 | -0.011 | 0.991 |  | -29.082 | 62.653 | -0.464 | 0.643 |
| **Pilis-Visegrád Mountains** | -0.041 | 0.341 | -0.119 | 0.905 |  | -4.413 | 57.697 | -0.076 | 0.939 |
| **Season (winter)** | 0.086 | 0.361 | 0.237 | 0.813 |  | 5.788 | 61.104 | 0.095 | 0.925 |
| **Jyvaskyla*winter** | 0.316 | 0.487 | 0.648 | 0.517 |  | -77.368 | 82.452 | -0.938 | 0.349 |
| **Turku*winter** | 0.511 | 0.527 | 0.969 | 0.333 |  | -0.913 | 89.137 | -0.010 | 0.992 |
| **Tartu*winter** | 1.069 | 0.629 | 1.700 | 0.091 |  | 36.466 | 106.407 | 0.343 | 0.732 |
| **Lund*winter** | 0.113 | 0.500 | 0.267 | 0.789 |  | 114.140 | 84.636 | 1.349 | 0.179 |
| **Pilis-Visegrád Mountains *winter** | 0.203 | 0.497 | 0.408 | 0.684 |  | 95.875 | 84.040 | 1.141 | 0.255 |

* La Hiruela and Westerheide populations (N = 31) are excluded from this model as those populations where only recorded during summer season. Oulu as the northernmost population is used as a reference to other populations and season (Summer) as a reference to season (Winter). 95 % confidence interval.

**B)**

| **Shannon** | **SumSq.** | **df** | **F value** | **P** | **Chao1** | **SumSq.** | **df** | **F value** | **P** |
| --- | --- | --- | --- | --- | --- | --- | --- | --- | --- |
| **Population** | 1.013 | 5 | 0.152 | 0.979 |  | 124 544 | 5 | 0.651 | 0.661 |
| **Season** | 0.075 | 1 | 0.056 | 0.813 |  | 343 | 1 | 0.009 | 0.925 |
| **Population*Season** | 4.727 | 5 | 0.708 | 0.618 |  | 290 113 | 5 | 1.517 | 0.185 |

* La Hiruela and Westerheide populations (N = 31) are excluded from this model as those populations where only recorded during summer season.

### SI 6. Permutational analysis of variance to measure which factors contribute to the differences in gut microbiome beta diversity.

| ***All data*** |  |  |  |  |  |
| --- | --- | --- | --- | --- | --- |
|  | **df** | **SumOfSqs** | **R2** | **F** | **P** |
| **Population** | 5 | 2.349 | 0.021 | 1.010 | 0.397 |
| **Season** | 1 | 0.592 | 0.005 | 1.272 | ***0.034*** |
| **Residual** | 239 | 111.218 | 0.974 |  |  |
| **Latitude** | 1 | 0.485 | 0.005 | 1.044 | 0.296 |
| **Body condition** | 1 | 0.478 | 0.005 | 1.031 | 0.353 |
| ****Weight*** | *1* | *0.478* | *0.005* | *1.030* | *0.355* |
| **Habitat** | 1 | 0.451 | 0.004 | 0.971 | 0.566 |
| **Temperature** | 1 | 0.629 | 0.006 | 1.355 | ***0.012*** |
| **Rainfall** | 1 | 0.385 | 0.004 | 0.829 | 0.961 |
| **Sex** | 1 | 0.456 | 0.005 | 0.983 | 0.486 |
| **Residual** | 212 | 98.435 | 0.972 |  |  |
| ***Winter*** |  |  |  |  |  |
| **Latitude** | 1 | 0.555 | 0.012 | 1.195 | 0.062 |
| **Body condition** | 1 | 0.420 | 0.009 | 0.905 | 0.809 |
| ****Weight*** | *1* | *0.420* | *0.009* | *0.905* | *0.810* |
| **Habitat** | 1 | 0.426 | 0.009 | 0.919 | 0.744 |
| **Temperature** | 1 | 0.428 | 0.009 | 0.923 | 0.753 |
| **Rainfall** | 1 | 0.408 | 0.009 | 0.878 | 0.889 |
| **Sex** | 1 | 0.506 | 0.011 | 1.091 | 0.168 |
| **Diet** | 1 | 0.511 | 0.011 | 1.100 | 0.164 |
| **Residual** | 92 | 42.715 | 0.929 |  |  |
| ***Summer*** |  |  |  |  |  |
| **Latitude** | 1 | 0.556 | 0.009 | 1.198 | 0.076 |
| **Body condition** | 1 | 0.430 | 0.007 | 0.926 | 0.705 |
| ****Weight*** | *1* | *0.436* | *0.006* | *0.941* | *0.679* |
| **Habitat** | 1 | 0.446 | 0.008 | 0.962 | 0.564 |
| **Temperature** | 1 | 0.539 | 0.009 | 1.162 | 0.093 |
| **Rainfall** | 1 | 0.448 | 0.008 | 0.965 | 0.552 |
| **Sex** | 1 | 0.439 | 0.007 | 0.946 | 0.604 |
| **Residual** | 122 | 56.617 | 0.952 |  |  |

*Each model was run first with body condition and then with weight to check consistency of results between both physiological metrics.
